# Cell migration driven by long-lived spatial memory

**DOI:** 10.1101/2021.01.05.425035

**Authors:** Joseph d’Alessandro, Alex Barbier-Chebbah, Victor Cellerin, Olivier Bénichou, René-Marc Mège, Raphaël Voituriez, Benoît Ladoux

**Affiliations:** Institut Jacques Monod (IJM), CNRS UMR 7592 et Université de Paris, 75013 Paris, France; Laboratoire de Physique Théorique de la Matière Condensée, CNRS / Sorbonne Université, 4 Place Jussieu, Paris, France; Laboratoire Jean Perrin and Laboratoire de Physique Théorique de la Matière Condensée, CNRS / Sorbonne Université, 4 Place Jussieu, Paris, France

## Abstract

Many living cells actively migrate in their environment to perform key biological functions – from unicellular organisms looking for food to single cells such as fibroblasts, leukocytes or cancer cells that can shape, patrol or invade tissues. Cell migration results from complex intracellular processes that enable cell self-propulsion ^1,2^, and has been shown to also integrate various chemical or physical extracellular signals ^3,4,5^. While it is established that cells can modify their environment by depositing biochemical signals or mechanically remodeling the extracellular matrix, the impact of such self-induced environmental perturbations on cell trajectories at various scales remains unexplored. Here, we show that cells remember their path: by confining cells on 1D and 2D micropatterned surfaces, we demonstrate that motile cells leave long-lived physicochemical footprints along their way, which determine their future path. On this basis, we argue that cell trajectories belong to the general class of self-interacting random walks, and show that self-interactions can rule large scale exploration by inducing long-lived ageing, subdiffusion and anomalous first-passage statistics. Altogether, our joint experimental and theoretical approach points to a generic coupling between motile cells and their environment, which endows cells with a spatial memory of their path and can dramatically change their space exploration.

---

Cell migration is essential for fundamental phases of development and adult life, including embryogenesis, wound healing, and inflammatory responses ^7^; it generically results from the active dynamics of its intracellular components – most prominently the cytoskeleton –, which generate propulsion forces and determine the cell front-rear polarity ^8,9^. The cytoskeleton spatio-temporal dynamics is controlled by complex regulatory networks ^10^, and can be characterized by both deterministic and stochastic components ^11,12,13,14^. The integration over time of this complex intracellular dynamics determines the large scale properties of cell trajectories, which can in turn be used as accessible read-outs to infer intracellular properties^2,15^, as well as cell interactions with the environment^5,16^ or with neighbouring cells ^17,18,19^. In vivo, cells interact with various extra-cellular environments with a broad range of biochemical and biomechanical properties ^8^. These interactions have been shown to be two-way: environmental cues directly affect cell shape, migration, and polarity^20,21,22^, and in turn, cells actively contribute to remodel their environment^6,23^. So far, however, both ways have been analysed independently, and the feed-back of cell-induced environmental remodeling on large scale properties of cell migration has remained unexplored.

To overcome the inherent complexity of the analysis of cell migration in 3D *in vivo* environments, the design of micropatterned surfaces has proven to be a powerful approach ^24,25,26^. In such *in vitro* set-ups, and especially in 1D settings, the reduced dimensionality of the cellular environment allows for an extensive quantitative analysis of the phase space roamed by migrating cells. In particular, such 1D assays have revealed striking deterministic features in cell motility patterns, while cell paths in higher dimensions remain seemingly random^27,14,28,6,19^. Moreover, many of the cell migration features on a 1D substrate can mimic cell behaviour in 3D matrix ^6^.

To dissect the mechanisms driving the spontaneous migration of living cells over a broad range of time scales, we followed single isolated MDCK epithelial cells (treated with mitomycin C to prevent cell division) on micro-contact-printed 1D linear tracks of fibronectin (Figure 1a-b). This set-up allowed us to track cells from their early spreading phase and on over long time scales (48 − 96 h) using video-microscopy (Figure 1b-c). By detecting the cell edges, we could reconstruct the trajectories of single cells in absence of cell-cell contact interactions. We observed two main behaviours: a first population of cells exhibited static spreading, and extended slowly their two ends in opposite directions without net displacement of their centre-of-mass (Figure 1d and Supplementary Video 1), while a second population displayed strikingly regular oscillatory trajectories, with an amplitude that could significantly exceed the cell size (Figure 1e and Supplementary Video 2). While oscillatory patterns in cell migration ^29,30^, and more generally in cell dynamics ^31,32,33^, have been reported for various cell types, and could be attributed to different intracellular processes, we argue below that the oscillations that we observe have so far unrevealed features, which we show originate from a so far unreported mechanism. Both observed behaviours were approximately equally distributed over the cell population, whereas behaviours that did not fall in these two classes – akin to persistent random motion – remained negligible. These observations were robust upon varying the width of the track over a range comparable to a single cell size, *W* = 10, 20, 50 µm (Figure 1f and Supplementary Figure 1a–c). To analyse the dynamics of front back cell polarity in these motility patterns, we used as a proxy for cell polarisation the concentration profile of a fluorescent biosensor (p21-activated kinase binding domain, PBD) of active Rac1 and Cdc42^24^, which are two well known activators of actin protrusions; as expected, moving cells displayed an increased PBD signal at the front, and lowered at the back, indicating their polarity. Two distinct phenotypes of polarisation, corresponding to the two observed dynamic behaviours clearly emerged from observations. On the one hand, static spreading cells were usually extended (up to *>* 100 µm) and characterized by a symmetric PBD profile with active poles at each of the two cell ends (see Supplementary Figure 1g and Supplementary Video 3). On the other hand, oscillating cells displayed “run” phases characterized by reduced cell length (∼ 20 µm), clear front back polarity, and roughly constant high speed (often faster than 100 µm.h^−1^, see Figure 1f-g, Supplementary Figure 1 and Supplementary Video 3-4). These run phases were interrupted by phases of polarity reversal, where the cell phenotype was transiently comparable to that of static spreading cells.

**Figure 1:**
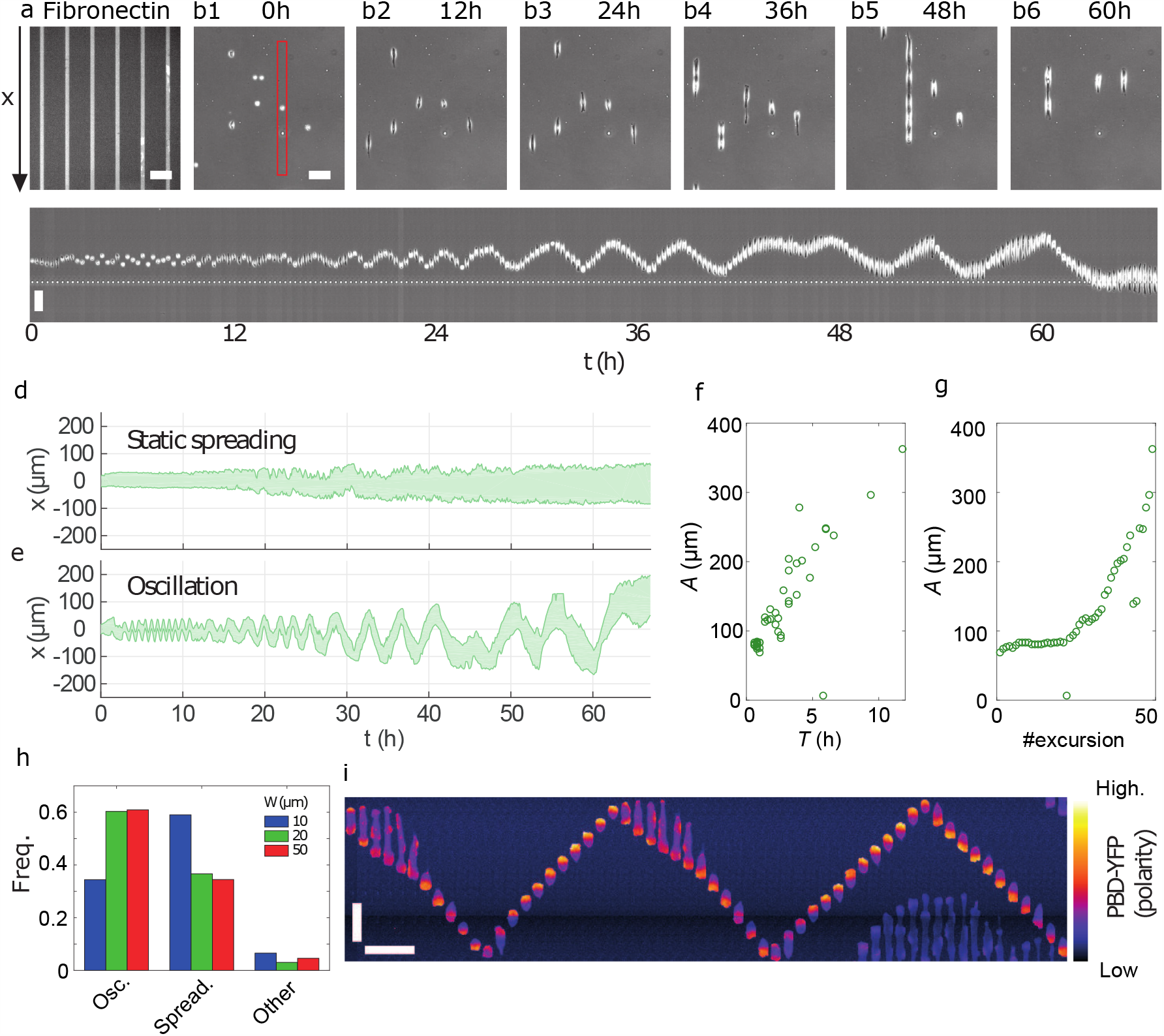
Isolated cells exhibit regular oscillations. ***a***. *Fluorescent tracks of fibronectin of width W* = 20 µm.. ***b1–b6***. *Snapshots of MDCK cells observed using phase-contrast imaging*. ***c***. *Kymograph of a single MDCK cell (red frame in* ***b1****) showing oscillations*. ***d-e***. *Typical kymographs of a statically spreading (****d****) and an oscillating (****e****) cell plated on lines with W* = 20 µm. ***f-g***. *Amplitude of the oscillations measured in panel* ***e*** *as a function of their period (****f****) and as a time series*. ***h***. *Frequency of the various behaviours (Oscillating, Spreading and Other) of isolated MDCK cells on lines of different widths. n* = 61, 131, *and* 87 *trajectories for W* = 10, 20 *and* 50 µm *from 6 independent experiments (2 per track width)*. ***i***. *Kymograph of a MDCK cell expressing PBD-YFP, a reporter of Rac1/Cdc42 activation, hence of the cytoskeleton polarity. Individual frames are separated by* 10 min. *All scale bars*, 100 µm.

We next focused on the oscillatory motility patterns, and characterized quantitatively their striking regularity. Despite some heterogeneity within the population, pointing to single-cell-specific properties, kymographs consistently displayed sawtooth-like patterns (Supplementary Figure 1). Both the period and amplitude increased in time, starting from very short (*A* ∼ 20 µm, *T <* 1 h) scales and reaching very high values up to *A* = 500 µm, *T* = 20 h (Supplementary Figure 2). Strikingly, within single trajectories the *A/T* ratio remained close to constant over this very broad range of values of *A* and *T*. This was consistent with cells running at roughly constant speed and bouncing between two virtual reflecting walls imposing polarity reversals, which would slowly move apart. Based on this observation, and because in our set-up external cues imposing such dynamics could be excluded, we hypothesized that the observed polarity reversals were induced by interactions of cells with their own footprints, and not caused only by an autonomous intracellular clock. More precisely, we conjectured that cells modify the physicochemical properties of their local environment, thereby leaving long-lived footprints along their path. In turn, footprints were assumed to induce local polarity signals that favor cell polarisation pointing towards previously visited areas, and away from unvisited areas, and thus to effectively attract cells, thereby slowing down the large scale spreading of trajectories. In this scenario, cells would therefore run persistently while within the previously visited domain, and reverse their polarity when reaching an edge, thereby incrementally extending the visited domain by overshooting the edge.

To challenge this hypothesis, we prepared ‘conditioned substrates’ on which a first batch of cells plated at high density was left migrating freely; we thereby expected the substrate to be fully covered by cellular footprints. After removing this first batch of cells, we plated a new batch of isolated cells on these conditioned substrates (Figure 2a), and performed the same analysis as in the control set-up (ie on substrates that were not conditioned by a first batch of cells). As compared to the control case characterized by slowly spreading oscillatory trajectories, cells on conditioned substrate displayed strikingly different migration patterns, with a drastically increased net displacement and a significantly larger persistence time, while only few oscillatory patterns could be observed (Figure 2b, c, e and Supplementary Video 5). In addition, cell spreading at early times and cell instantaneous speed were found to be larger on conditioned substrates (Supplementary Figure 3), indicating that cellular footprints facilitate adhesion and migration. This was further confirmed ^34^ by measuring the forces exerted by the cells on the substrate using traction force microscopy (TFM, Figure 2f)^29^. In this 1D setting, the strength of the coupling between the cell and the substrate may be simply assessed by computing the maximal cell tension, which is obtained by integrating the 1D traction force profile along the line pattern (Figure 2g-h). We thus concluded that cells on conditioned substrates are able to exert traction forces twice as large as cells on control substrates (Figure 2i). Importantly, these results are not limited to the 1D linear geometry. We repeated the motility assay on homogeneously coated surfaces and characterised the dynamics of free 2-dimensionnal cell trajectories: comparing substrates with and without conditioning by a first batch of cells, we observed a 4-fold increase of the effective diffusion constant on conditioned substrates as compared to control substrates (Figure 2d-e). Finally, we confirmed our observations by analysing another cell type: Isolated Caco2 – colorectal cancer – cells also exhibit oscillations on control linear patterns, while on substrates conditioned by a first batch of Caco2 cells they move persistently (Figure 2e and Supplementary Figure 11). Altogether, these results support our hypothesis of a generic phenomenon of cellular footprint deposition, which deeply impacts cell trajectories by restricting them to already visited areas.

**Figure 2:**
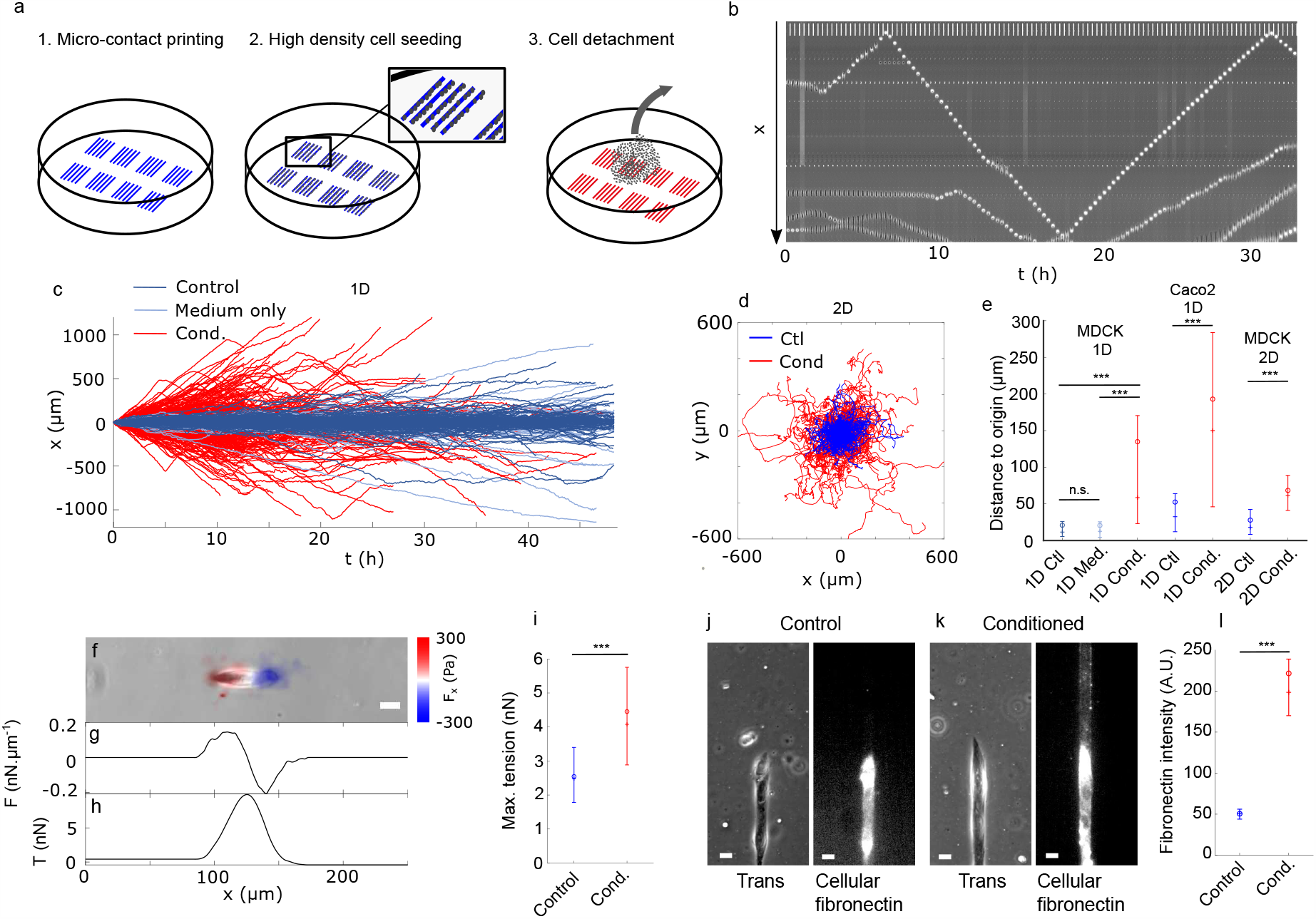
Cells deposit a footprint on their tracks. ***a***. *Principle of the substrate conditioning. Linear tracks were micro-contact printed (1), then a first layer of cells was plated at high density to recover all the surface (2) before being detached (3)*. ***b***. *Kymograph of a cell moving along a conditioned* 20 µm *track with high persistence. Scale bar (repeated vertical line)* 100 µm. ***c***. *Trajectories of cells on control (Ctl*: *substrate kept in PBS, Medium only: substrate kept in DMEM during the same time as the conditioning) or conditioned susbtrates as a functions of time. This shows high persistence on conditioned susbtrates. Only trajectories of at least* 10 h *duration are shown*. ***d***. *Trajectories of single cells plated on 2D surfaces on control (Ctl, blue) or conditioned (Cond*., *red) substrates*. ***e***. *Distance of the cell to its original position after* 16 h *on control and conditioned substrates in 1D and 2D. Differences were assessed using the 2-sample Kolmogorov-Smirnov test, n*.*s*.: *non-significant (p >* 0.1), ***: *p <* 0.001. *n* = 355, 238, 429, 246, 216, 192, *and* 194 *trajectories from 3 (MDCK 1D, Ctl / Medium / Cond*.*), 3 (Caco2 1D, Ctl / Cond*.*) and 2 (MDCK 2D, Ctl / Cond*.*) independent experiments respectively*. ***f***. *Phase-contrast image of a MDCK cell on a soft PDMS substrate, overlaid with traction stress. Scale bar* 20 µm. ***g***. *Traction force profile along the cell, integrated over the line width*. ***h***. *Tension T within the cell obtained by integrating the traction force profile along the x-axis*. ***i***. *Maximal (peak) tension of cells on control (blue) or conditioned (red) linear substrates. Difference between n* = 26 *and* 31 *cells was tested using the 2 sample Kolmogorov-Smirnov test*, ***: *p* = 3 × 10^−5^. ***j-k***. *Phase-contrast and immuno-staining of cellular fibronectin on a control (e) and a conditioned (f) line. In both cases, there is a cell only on the bottom part. Scale bars* 20 µm. ***l***. *Cellular fibronectin intensity in control (blue) and conditioned (red) lines devoid of cells. Difference between n* = 246 *and* 66 *independent lines for control and conditioned substrates respectively was tested using the 2 sample Kolmogorov-Smirnov test*, ***: *p <* 1 × 10^−3^. *The plots in panels* ***e, i*** *and* ***l*** *display the mean (o), median (+) and first and third quartiles (error bars) of the distributions*.

**Figure 3:**
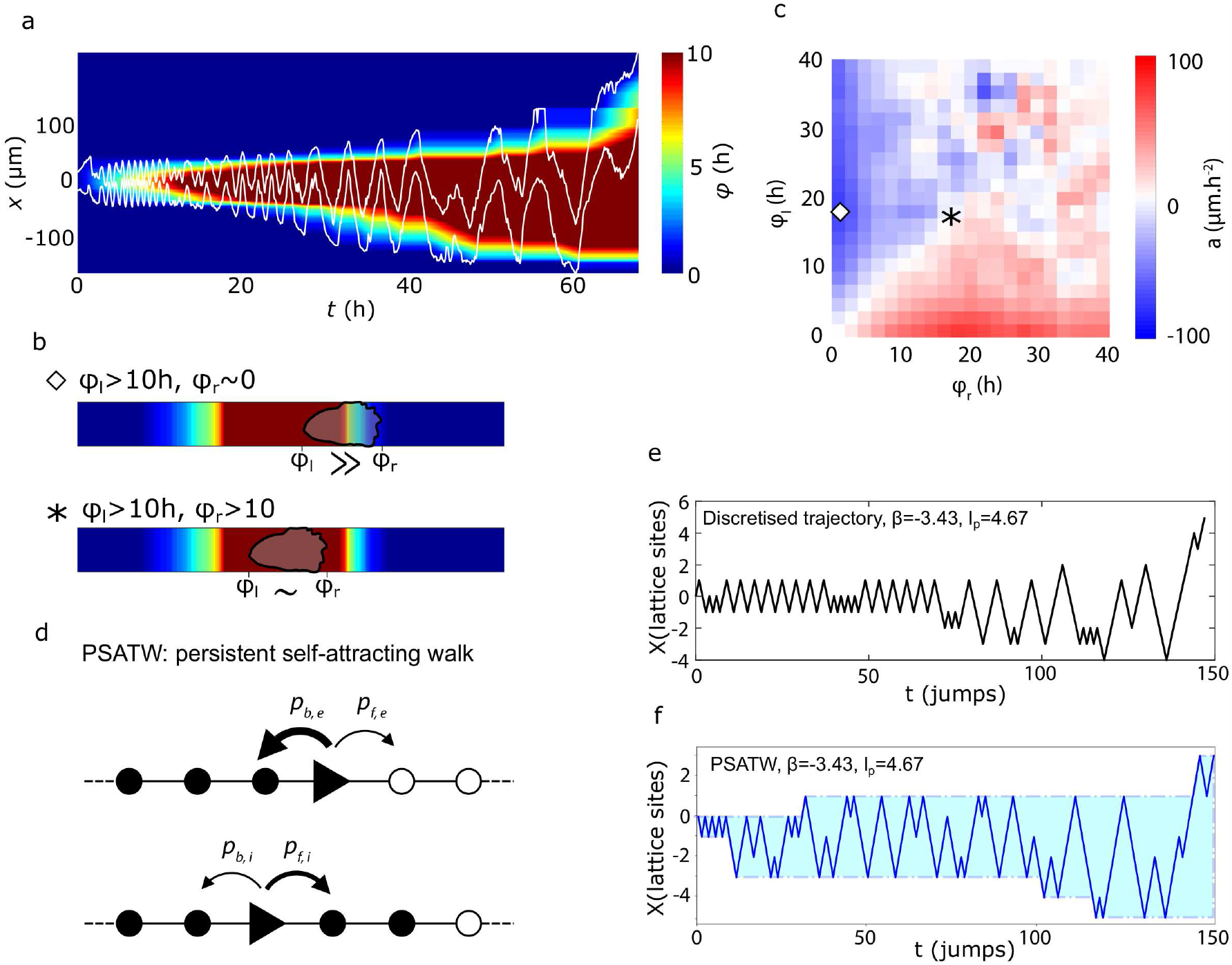
The trajectories of isolated cells have the characteristics of self-attracting walks. ***a***. *Kymograph of an isolated cell on a W* = 20 µm *track, overlaid with its footprint field φ*(*x, t*) *defined as the cumulative time spent on a given position*. ***b***. *Sketch of the φ*_*l*_ *and φ*_*r*_ *measurements. Top: the cell sits on the right edge of the footprint, its right edge being outside with φ*_*r*_ ∼ 0 *while its left edge is within, φ*_*l*_ *>* 10 h. *Bottom: the cell is completely within the footprint, with comparable high values of φ*_*l*_ *and φ*_*r*_. *The symbols are reported in panel* ***c*** *to show where each situations sits in the φ*_*l*_ − *φ*_*r*_ *space*. ***c***. *Average acceleration of isolated cells on* 20 µm *tracks as a function of the values of φ at both cell ends*. ***d***. *Sketch of the persistent self-attracting walk model. Whether it is on the edge of its footprint (top) or in its interior, the walker have different probabilities to jump in the same direction as before or to turn back, set by two parameters k and β*. ***e***. *Discretised experimental trajectory of an oscillating cell (same cell as panel a) allowing to measure the reversal statistics within and on the edge of the span*. ***f***. *Simulated trajectories of an agent following the PSATW dynamics with the same parameters as infered in panel* ***e***.

Next, to substantiate our hypothesis, we sought to identify chemical components of the cellular footprints. It is known that cells are able to assemble or produce extra-cellular matrix proteins, and in particular fibronectin ^35^, which thus appeared as a natural candidate. In order to distinguish cell-produced fibronectin from pre-coated fibronectin, after fixation samples were stained using an antibody directed specifically to the fibronectin produced by the cells themselves^36^. In the 1D set-up, we observed that cellular fibronectin localised only in limited areas previously visited by cells, whereas no deposited fibronectin could be detected in areas that remained unexplored by cells (Figure 2j). Consistently, on conditioned substrates tracks were found to be fully covered with cellular fibronectin, with a higher density along the tracks edges (Figure 2k-l). We also added marked plasma fibronectin in the medium, thus making it available for cells to capture, assemble and deposit along their path ^66^. This strategy yielded similar results as the staining of cellular fibronectin, showing that cells are also able to deposit fibronectin that is available in the medium (Supplementary Figure 10). These results suggest that ECM deposition participates to memory effects in cell migration. However, we cannot exclude that other molecular or supra molecular components, such as exosomes ^37^ or cell fragments ^38^ could be released as well, and even the sole remodelling of the pre-existing extra cellular matrix could be invoked; a complete description of cellular footprints would go far beyond this work. However, our results provide a direct evidence that cells indeed leave long lived chemical footprints – made at least of fibronectin –, which, as we showed, can deeply modify cell motion at later times.

To characterize the impact of cellular footprints on cell dynamics at the cell scale, we developed a kinematic approach based on the analysis of 1D cell trajectories. As a proxy for any potential deposited signal, we defined a footprint field *φ*(*x, t*) as the cumulative time spent by a cell on a given location *x* before time *t* (Figure 3a). We next analysed the correlations between the acceleration *a* of the cell centre-of-mass, and the *φ* values measured at the left (*φ*_*l*_) and right (*φ*_*r*_) ends of the cell (Figure 3b-c and Supplementary Figure 4–7). For a cell moving within the previously visited domain, the footprint field probed by the cell is roughly uniform (*φ*_*l*_ ∼ *φ*_*r*_ » 0), and we observed no significant variation of the cell migration speed (*a* ∼ 0). In contrast, for a cell reaching for example the right (resp. left) end of the visited domain the local footprint field probed by the cell is very asymmetric with *φ*_*l*_ » *φ*_*r*_ (resp. *φ*_*l*_ « *φ*_*r*_), and we observed a significant average acceleration inward the visited domain, (*a*) ≃ −100 µm.h^−2^ (resp. *a* ∼ +100 µm.h^−2^). This clear correlation indicates that cell polarity is governed by local gradients *δφ* = *φ*_*r*_−*φ*_*l*_ of the footprint field, and substantiates our earlier hypothesis that cellular footprints impact on cell trajectories.

Our experimental results show that cells, by leaving chemical footprints along their way, are endowed with a spatial memory of their path. Their theoretical analysis therefore calls for a framework that goes beyond the classical models invoked in the literature, which are for most of them amenable to markovian, and therefore memoryless descriptions^11,12,13,16,18^, with the exception of ^40,41^. Our observations led us to argue that cell trajectories naturally fall in the class of self interacting random walks, which can be broadly defined as the class of random walks that interact (attractively or repulsively) with the full territory explored until time *t*^42,43,44,45,46,47,48,49^. This class comprises in particular the self-avoiding random walk, which has played a crucial role in physics^50^, and has applications in the modelling of trajectories of living organisms ^51,52,53^. By construction, self-attracting random walks are endowed with long range memory effects, which makes their analytical study notoriously difficult.

A generic example of self interacting random walk is given by the so–called Self-Attracting Walk (SATW). It can be defined on a 1D lattice as a discrete time random walk whose jump probability to a neighboring site *i* is assumed to be proportional to exp(−*βf* (*n*_*i*_)), where *n*_*i*_, defined as the number of times site *i* has been visited by the random walker up to *t*, is akin to the footprint field *φ* defined above. Upon varying the parameter *β <* 0 and the increasing function *f* (case of self attraction), this model is known to display a broad range of behaviours, from everlasting trapping on a few sites to large scale diffusion^46^. More specifically we used the SATW to build explicitly a minimal model of cell dynamics that recapitulates our main observations on migrating cells. For the sake of simplicity, we took *f* (*n*_*i*_ = 0) = 0 and *f* (*n*_*i*_ *>* 0) = 1, which amounts to assuming that the deposited signal that defines cellular footprints is bounded. Next, we extended the SATW model to take into account cell persistence. The persistent self-attracting walk (PSATW) can be defined as follows in 1D. When the walker is on a site *i* within the visited domain – ie surrounded by sites that have already been visited, *n*_*i*−1_, *n*_*i*+1_ *>* 0 – it performs a classical persistent random random walk: it changes direction with probability 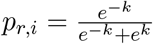, and reproduces its previous step with probability 1 − *p*_*r,i*_, where *k >* 0 is a parameter that controls the cell persistence length *l*_*p*_ = *e*^2*k*^ (the persistence time *t*_*p*_ is defined identically by setting cell speed to 1). When the walker is at an edge of the domain (eg *n*_*i*+1_ *>* 0), it experiences a local bias inward the visited domain parametrized by *β <* 0 and the probability to change direction can be written 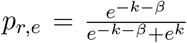, while the probability to reproduce the previous step is 1 − *p*_*r,e*_ (Figure 3d). With this definition, a typical PSATW trajectory with *k >* 0 and *β <* 0 shows noisy oscillatory patterns, with an amplitude that slowly increases over time, which qualitatively reproduce experimental observations (Figure 3e-f). More quantitatively, we found that the experimental trajectories, after adequate discretization, could be well fitted by adjusting the *k, β* parameters, with an inherent cell to cell variability (Supplementary Figure 8). Of note, the fitted *k* values, that control the intrinsic cell persistence length, were comparable on conditioned and control substrates, while the parameter *β*, which controls the cell response to the footprint field, was significantly different in both conditions (Supplementary Figure 9). This finally shows that PSATWs provide a minimal model of the self interacting random walk class, which reproduces the observed migration patterns.

Last, we show both theoretically and experimentally that the reported interaction of cells with their footprint, which endows cells with a memory of their path, has important consequences on space exploration properties of cell trajectories. (i) First, the time dependent increments defined by *I*(*T, t*) ≡ ⟨[*x*(*t* + *T*) − *x*(*T*)]^2^), which quantify the spreading speed of trajectories, are found to depend on the measurement time *T* at all time scales, ie to display ageing, in both 1D and 2D set-ups and in agreement with the 1D PSATW model and the 2D SATW model (Figure 4a,b,d,e). Conversely, we observed that aging of the increments was negligible on conditioned substrate (Figure 4c,f), further confirming our findings that memory effects where induced by cellular footprints. This is a direct consequence of the increase over time of the span of the visited territory. In 1D, at short measurement times *T* « *t, t*_*p*_, the increments dynamics is governed by interactions events with the edges of the visited domain, which slow down spreading and lead to a diffusive behaviour *I*(*T, t*) ∼ *t* for both *t < t*_*p*_ and *t > t*_*p*_. For *T* » *t, t*_*p*_, edge effects become negligible and one recovers the classical dynamics of persistent random walks, which crosses over from a ballistic (*I*(*T, t*) ∼ *t*^2^) to a diffusive (*I*(*T, t*) ∼ *t*) regime. In 2D, the observed persistence length is comparable to the cell scale and can be neglected; one can thus use a classical SATW model. This model was shown to lead to normal diffusion in the *T* » *t* regime, and to subdiffusion ^46,47^ *I*(*T, t*) ∼ *t*^2*/*3^ in the *T* « *t* regime, which is consistent with our observations, even if experimental data do not allow to determine quantitatively the exponent. This subdiffusive regime shows that memory effects can have drastic consequences on space exploration, by changing the very dimension of trajectories, which qualitatively become more compact. (ii) Second, we argue theoretically that such ageing dynamics has important consequences on first-passage time statistics ^54,55^, which is a key observable to quantify the efficiency of migratory patterns to find “target” sites of interest in space^56,57^. In the simplest theoretical setting of a single target located in infinite space, first-passage statistics to the target are conveniently parametrized by the persistence exponent *θ*, which defines the long time asymptotics of the survival probability of the target *S*(*t*) ∝ *t*^−*θ* 58^. For a broad class of random walks, which do not display ageing, the persistence exponent takes the remarkable universal value *θ* = 1*/*2^58^. Strikingly, our results show that memory effects in the PSATW model in 1D lead to non trivial values of *θ* ^55^, which are controlled by the parameter 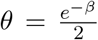, Figure 4c). Of note, one has *θ >* 1*/*2, indicating that memory effects increase the relative weight of shorter trajectories and thus favor local space exploration, as compared to memoryless random walks with *θ* = 1*/*2. First-passage statistics are thus deeply impacted by memory effects in the PSATW model.

**Figure 4:**
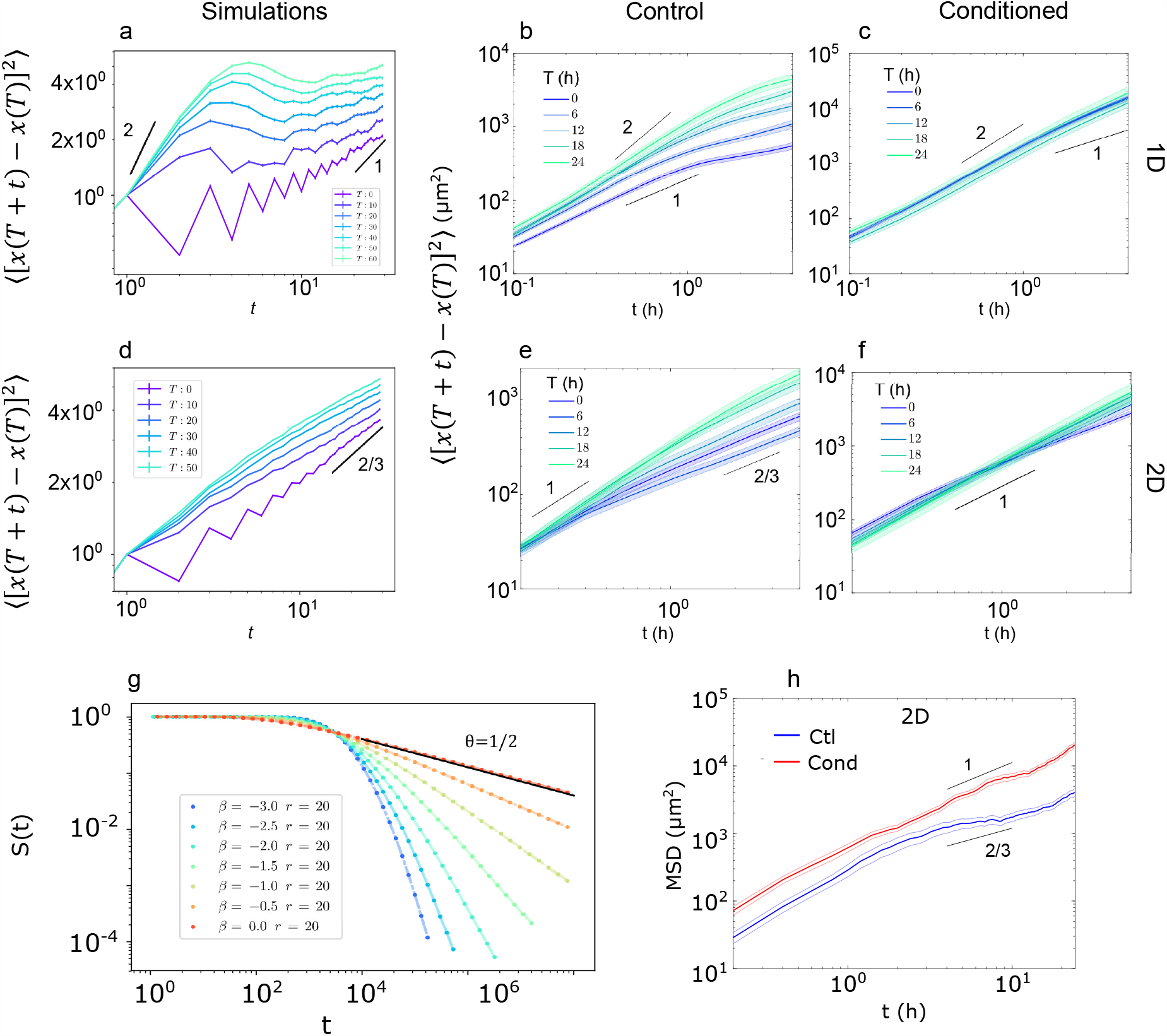
The PSATW model predicts altered cell trajectory statistics and a better exploration of space in both 1D and 2D. ***a–f***. *Increments of the mean square displacements in 1D (a–c) or 2D (d–f) settings. Data from simulations (a,d) and experiments on control (b,e) and conditioned (c,f) substrates. Lines are guides for the eye showing the various exponents predicted by the theory. In panels* ***b, c, e***, *and* ***f*** *the error bars show the S*.*E*.*M. for n >* 100 *trajectories from 3 (1D) and 2 (2D) independent experiments*. ***b***. *Survival probability S*(*t*) *of a target located at a distance r of the origin in 1D as a function of time. The long-time scaling θ exponent decreases as* 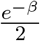. ***d***. *Mean square displacement of isolated cells moving on 2D surfaces in logarithmic scale. Control (Ctl, blue) and conditioned (Cond*., *red) substrates. The error bars show the S*.*E*.*M. for n* = 203 *and* 172 *trajectories from 2 independent experiments for control and contitioned substrates respectively*.

Moving cells interpret multiple physical ECM parameters in parallel and translate them into an integrated response, which determines cell shape, polarity and migration ^59^. From this well-established principle, our results provide further steps in our understanding of cell-matrix interactions. We indeed demonstrate that reciprocal dynamic adaptation between cell and its external environment is crucial to determine cell migration principles. Cells not only respond and adapt to their environment, but they are also able to build their own road while advancing. Here we show how cells keep track of their previous locations by depositing a footprint on their path, and, in turn, how this footprint determines dynamical properties of cell migration. Our discovery of self-attraction mechanisms during cell migration manifests through the emergence of oscillatory modes when cell motion is confined on a 1D line and peculiar sub-diffusive trajectories over 2D surfaces.

In fact, we anticipate that these feedback mechanisms between cell migration and substrate modifications could play an important role in many other situations where it was not suspected. In vivo, motile cells integrate various inputs from the extra-cellular matrix, and, then, adjust their migration mode ^60^. Gagné *et al*. ^61^ showed that ILK potentiates intestinal epithelial cell spreading and migration by allowing fibronectin fibrillogenesis. The assembly of new roads built up by the cells themselves can be also crucial in the context of collective cell migration where the remodeling of ECM by leader cells can provide a path for follower cells^62^. Such retroactions between intra- and extra-cellular dynamics could thus be a more generic feature than it is currently considered and previous works on cell migration might be revisited in the light of our findings. In the present case, the self-attraction mechanism described here has three main consequences that might be of biological significance. First, it generates loose self-confinement, which at first sight prevents efficient migration over long distances. In return, this localisation ensures a much better exploration of space, so that no unexplored ‘hole’ is left behind, which might be of importance for cells that need to patrol a zone. Finally, it confers ageing on cell trajectories, which might be crucial although overlooked in cell migration experiments: it means that the movement properties at a given time can depend strongly on the interval between the time at which cells are deposited and the time at which measurements start. Two questions remain open: the nature of the footprint and the physical mechanism by which the cell is attracted back. Several works have shown that cells are able to produce and to remodel their extra-cellular matrix ^35,61,63,64,65,66^. In this study, we have shown that cells actually deposit fibronectin in the form of small puncta but other ECM components may play a role. In addition, other candidates have been evidenced as self-attraction media, notably exosomes ^37^. The mechanism of repolarisation at the footprint edge also remains unclear, although its details could be of importance for the overall dynamics: the cell-substrate system needs to be close to a specific operation point, such that the footprint is deposited efficiently enough for the cell to sense it, but not too fast so the cell reverses its polarity before it has built a strong enough footprint at its front to keep moving ahead.

Our findings provide a novel framework to understand the intimate relationship between cell and ECM remodelling. Even tough our study focuses on single cell behavior, we anticipate that it could play a role in collective cell dynamics. In living tissues involving cell populations, either sparse or dense, either homogeneous or heterogeneous, reinforced motion could manifest through a broad variety of consequences. For instance, Attieh *et al*. ^65^ showed that cancer associated fibroblasts could open the way to cancer cells by assembling fibronectin fibrils along collagen fibres. There is no doubt that such amazing collective effects arise when several self-attracting walks interact, or if the self-attracting field can be degraded with time. Future studies may try to introduce those levels of complexity and analyse the role they play in various physiological and pathological situations.

## Acknowledgements

We thank the group members from “Cell adhesion and migration” team for helpful discussions. This work was supported by the LABEX Who Am I? (ANR-11-LABX-0071), the Ligue Contre le Cancer (Equipe labellisée 2019), and the Agence Nationale de la Recherche (‘POLCAM’ (ANR-17-CE13-0013 and ‘MechanoAdipo’ ANR-17-CE13-0012). We acknowledge the ImagoSeine core facility of the IJM, member of IBiSA and France-BioImaging (ANR-10-INBS-04) infrastructures.

## Authors’ contributions

JdA, BL and RV designed the research. JdA performed the experiments and analysed the experimental data, except TFM experiments, which were performed and analysed by VC. ABC, OB and RV designed the model. ABC performed the numerical simulations and analysed the simulation data. JdA, ABC, BL and RV wrote the manuscript. All authors commented on the manuscript and agreed on its final version.

## Data and code availability

Data and analysis codes supporting this paper are available upon request for result reproduction purposes.

