## Supplementary Information file for "Cell migration driven by long-lived spatial memory"

### Cell migration driven by long-lived spatial memory
### Supplementary material

Joseph d'Alessandro,[1,*] Alex Barbier-Chebbah,[2] Victor Cellerin,[1] Olivier Benichou,[2] René-Marc Mège,[1] Raphaël Voituriez,[2,3,†] and Benoit Ladoux[1,‡]

[1]*Institut Jacques Monod, CNRS UMR 7592, University Paris Diderot, Paris 75013, France*
[2]*Laboratoire de Physique Théorique de la Matière Condensée, UMR 7600 CNRS, Sorbonne Université, 75005 Paris, France*
[3]*Laboratoire Jean Perrin, UMR 8237 CNRS, Sorbonne Université, 75005 Paris, France*

#### I. Experimental procedures

##### A. Cell culture

We used MDCK wild-type, MDCK histon-GFP, MDCK PBD-YFP (gift from F. Martin-Belmonte lab) and Caco2 cells. The cells were cultured in DMEM GlutaMAX high-glucose (Gibco, Waltham, MA) supplemented with 10% fœtal bovine serum (FBS, BioWest, Nuaillé, France) for MDCK lines, and 20% FBS + 1% penicillin-streptomycin for Caco2. Prior to experiments, the cells were treated with $10\,\mu\mathrm{g.mL}$ of Mitomycin C added in the medium for $1\,\mathrm{h}$, then rinsed before subsequent detachment and seeding on the experimental samples.

##### B. Micro-contact printing

All micropatterns were prepared using standard micro-contact printing on PDMS, as described previously [1]. The substrates used were non-culture treated plastic dishes (Greiner Bio-One, Kremsmünster, Austria) for experiments in Figure 1, glass coverslips (Menzel-Gläser) for PBD-YFP experiments and 6-well plates for all other experiments. The substrates were first covered with a thin layer of poly-dimethyl-syloxane (PDMS, Sylgard, Dow Corning, Midland, MI)) using a spin-coater (or spread by manually, slowly rotating plates) and crosslinked at $80°\mathrm{C}$ for $2\,\mathrm{h}$. PDMS stamps were prepared by pouring PDMS on a mold featuring the patterns to be printed and crosslinked as described. After cooling down, a fibronectin solution was prepared by adding $50\,\mu\mathrm{g.mL}^{-1}$ of fibronectin and $25\,\mu\mathrm{g.mL}^{-1}$ of Cy3- or Cy5-labelled fibronectin into sterile milliQ water. The solution was then incubated on the stamps for $40\,\mathrm{min}$ at room temperature. Before stamping, the substrates were activated using UV-ozone for $10\,\mathrm{min}$ and the stamps were rinsed to remove excess fibronectin and dried using an air-gun. The stamps were briefly put in contact with the substrate's surface, then removed, and the substrates immerged in a $2\%$ pluronics F127 (Sigma-Aldrich, Saint Louis, MO) solution in PBS for $2\,\mathrm{h}$. Finally, the substrates were rinsed in PBS and sterilised under the UV lamp of a culture hood before use. The substrates for 2D migration were prepared similarly, using a large piece of flat PDMS as a stamp for homogeneous coating.

##### C. Substrate preparation for traction force microscopy

For traction-force microscopy (TFM) experiments, soft PDMS substrates were prepared and micro-contact printed as described previously [1]. Soft PDMS with Young's modulus of approximately $30\,\mathrm{kPa}$ was prepared by mixing components A:B of soft PDMS in a 5:6 weight:weight ratio, then poured on a 6-well plate and cured at $80°\mathrm{C}$ for $2\,\mathrm{h}$. Fluorescent beads (Fluorospheres, Invitrogen, Carlsbad, CA) were stuck on the surface after functionalisation with aminopropyl-triethoxy-silane (APTES, Sigma, Saint Louis, MO). Finally, the substrate was patterned by first using micro-contact printing on a poly-vinyl-alcohol (PVA, Sigma, Saint Louis, MO) membrane. Then the membrane was put in contact with the substrate for $30\,\mathrm{min}$, dissolved in water and rinsed before passivation with Pluronic F127 as above.

##### D. Conditioned substrates

To prepare conditioned substrates, substrates were first prepared as above. Conditioning cells were seeded at high density so as to cover all the available surface, rinsed after $45\,\mathrm{min}$ to prevent overcrowding and placed in the incubator. After $12\,\mathrm{h}$ of conditioning, the good covering of the substrate was checked under the microscope, and the culture medium was replaced with a $20\,\mathrm{mM}$ solution of EDTA in calcium- and magnesium-free PBS (Gibco, Waltham, MA). After $2\,\mathrm{h}$, the remaining cells were detached by flowing gently the solution using a pipette, and the substrate was rinsed with PBS.

---

[*] Electronic address:
[†] Electronic address:
[‡] Electronic address:

##### E. Cell deposition

Cells were detached using Trypsin-EDTA (Gibco, Waltham, MA), counted, centrifuged, resuspended in a 1:1 mix of full culture medium and low calcium DMEM to prevent too much cell clustering. Typically $10 - 20.10^3$ cells were seeded per well in control experiments on lines, while only $1.10^3$ were enough for experiments on conditioned substrates (even lower densities were used for experiments on 2D surfaces). Precise control of the cell density was necessary to ensure good statistics with a high number of cells remaining isolated for a long duration. After $45\,\mathrm{min}$, the sample was rinsed in PBS with great care to remove most floating cells without affecting recently adhered cells and $3\,\mathrm{mL}$ of full culture medium was added.

##### F. Live imaging

Live experiments were performed using a fully automated inverted microscope (Olympus, Japan) with computer-controlled temperature and CO2 level. At least one fluorescent image of the patterns was taken at the beginning of the experiment. Cells were imaged using phase-contrast imaging, with a time-lapse of 6 (1D), 10 (TFM) or 12 (2D) minutes between frames. For 2D experiments, histon-GFP expressing cells were used and imaged in fluorescence to make cell detection easier. For TFM experiments, the focus was done on the substrate's surface and fluorescent images of the beads were acquired at each frame. At the end of the experiment, the cells were detached using Sodium DodecylSulfate (SDS, Sigma, Saint Louis, MO) at 10% and an image of the beads was taken as a force-free reference. The imaging of PBD-YFP was done using an inverted microscope (Leica, Wetzlar, Germany) with a CSU-W1 confocal spinning-disk module (Nikon, Tokyo, Japan) and a 40X oil-immersion objective. The acquisition was done using Metamorph (Molecular Devices, San Jose, CA), at a 6 to 10 minutes acquisition rate. The focus was done on the basal plane of the cells and the microscope's hardware autofocus was used to ensure the absence of defocusing.

##### G. Labelled fibronectin deposition

To check for the deposition of fibronectin from the medium by the cells, $5\,\mu\mathrm{g.mL}^{-1}$ of Cy3- or Cy5-labeled fibronectin was added in the medium after the cell-rinsing step – choosing the colour in order to avoid confusion with the micro-contact-printed fibronectin. The cells were then left in the incubator for $48\,\mathrm{h}$ to allow them to deposit the labelled fibronectin along their path. The sample was imaged, either in PBS or mounted in Moewiol, after fixation and potential complementary immuno-staining.

##### H. Immuno-staining

The sample was fixed in 4% paraformaldehyde for $15\,\mathrm{min}$, then rinsed with PBS. The primary antibody (anti-[IST9] fibronectin, AbCam ab6328) was added at $1/100$ [4] and left at $4°\mathrm{C}$for $48\,\mathrm{h}$, then rinsed with PBS. The secondary antibodies were added at $1/250$ and left to incubate at room temperature for $2\,\mathrm{h}$, then rinsed with PBS. Finally, the sample was either placed in PBS (for 6-well plates) or mounted on a glass slide with Moewiol.

##### I. Image analysis – cell migration

The image treatment was performed using homemade ImageJ and Matlab (The Mathworks, Natick, MA) software. In brief, patterns were first detected using semi-automated methods, images were rotated so as to align all patterns on the same direction and positioning noise was removed if necessary using "Template Matching" ImageJ plugin. Cell contours were detected using the Edge detection function of ImageJ and thresholding with filters on the detected features' size and circularity, and the positions of both cell ends were recorded. For 2D experiments, the nucleus centre-of-mass was detected using a in-house program based on the Find Maxima function of ImageJ [2]. The cells were then tracked in time using the track.m (http://site.physics.georgetown.edu/matlab/) program in Matlab to reconstruct full cell trajectories. The trajectory data were organised as in [2] and adapted software was developed for all further analysis (footprint, velocity, auto-correlations, oscillation detection, MSDs...).

##### J. TFM analysis

The TFM data was analysed as previously described [1]. First, the displacement field of the beads was measured using particle-image velocimetry (PIV), using the matpiv (https://www.mn.uio.no/math/english/people/aca/jks/matpiv/) function in Matlab. Then the traction-stress (or maybe more accurately "traction-force-per-unit-area") field $\mathbf{f}(x, y, t)$ was computed using the FTTC [3] program in ImageJ
(https://sites.google.com/site/qingzongtseng/tfm), yielding For each time frame, the cell mask was extended so as to engulf all the corresponding force patch, and the force profile along the cell length was computed by averaging the forces within this mask along the line direction, $F = \langle f_x(x, y, t)\rangle_{y\in\mathrm{line}}$. The 1-dimensionnal tension within the cell (in the $x$-direction) is then simply the integral of this 1D force profile, $T(x, t) = \int_0^x -F(x, t)\mathrm{d}x$. As a good estimate for the maximal tension within the cell, we used half the integral of the asbolute value of the force profile long the cell. For each cell, the median total force was calculated over the experimental time.

##### K. Fluorescent fibronectin analysis

To measure the intensity of fluorescence of immuno-stained cellular fibronectin, we proceeded as follows. First, pictures were taken with the wide-field microscope to get quantitative data. The issue of this imaging technique is that the illumination field is very inhomogeneous. To remove this bias, we sought to measure locally the difference of fluorescence between the inside and the outside of the line patterns. To that end, we defined local ROIs of $130\,\mu\mathrm{m}$ length along the line and $60\,\mu\mathrm{m}$ width centered on the line center. We computed the intensity profile *in the region outside the line*, averaged over the ROIs *length* along the line. This profile was fitted with a linear function and the intensity profile corrected so that the intensity outside the line was fluctuating around 0. Finally, the intensity at a given location along the line was computed as the average of this corrected intensity over the line width. The location of cells on the lines was detected by simply thresholding the histon-GFP intensity. Then for all lines completely devoid of cells, we computed the median of the corrected intensity profile. For lines with cells, the corrected intensity was binned as a function of the distance to the closest cell nucleus, and the median was taken for each distance bin to show the decay of fluorescence away from cells.

##### L. Discretisation procedure

To analyse cell trajectories in the framework of the PSAW model, we discretised the experimental trajectories as follows. The centre-of-mass position $x$ was interpolated on a regular grid $X = \lfloor \frac{x}{L_{\mathrm{ref}}} + 0.5 \rfloor$ whose mesh size $L_{\mathrm{ref}}$ was defined as a cell-specific typical length. Then the time was interpolated on the irregular grid $(T_i)_{i\in\mathbb{N}}$ corresponding to hopping events $X(T_i) = X(T_i - 1) \pm 1$. Note that this way, time is irregularly interpolated. The distribution of jump times $(t_j^i) = (T_{i+1} - T_i)$ is bi-exponential with a mean of $\langle t_j \rangle \simeq 0.6h$, yielding discretised experimental trajectories of typically 100–150 time-steps (Figure 3f of the main text). Directly measuring the conditional frequencies of reversal $p_{r,i}$, and $p_{r,e}$, we computed one $(k, \beta)$ parameter set *per* cell by inverting the expressions of those probabilities in the PSAW model formulation.

##### M. MSD and ageing

The MSD plotted in Figure 4h is simply computed for each cell as $\|\mathbf{x}(t) - \mathbf{x}(0)\|$. For the ageing of the MSD, the trajectories were first cut in 12 h-long pieces starting at $T \in \{0, 6, 12, 18, 24\}$ h. The MSD was computed as $MSD(T,t) = \langle \|\mathbf{x}(t + t_0) - \mathbf{x}(t_0)\| \rangle_{t_0}$, where the angle brackets denote a sliding average over $t_0 \in [T; T + 12\,\mathrm{h}]$ to ensure better statistics, and the curve was cut at $12/3 = 4\,\mathrm{h}$ for the same reason.

#### II. Supplementary information – experiments

The Supplementary Figures 1 shows a substantial subset of the experimental cell trajectories on tracks of three different widths $W = 10, 20, 50\,\mu\mathrm{m}$ (here, only trajectories of $> 40\,\mathrm{h}$ have been retained). From those plots, one can appreciate at the same time both the generic aspect of the oscillation phenomenon and its variability. In what follows we tackle various aspects of this phenomenon.

##### A. Cell length and polarisation

By comparing the cell length in the two stereotypical examples shown in Figure 1 (static and oscillating cell), it appears that they share the same long time spreading dynamics – which might be largely due to mitomycin C treatment. The difference in the two behaviours should then arise simply from the capacity of cells to polarise strongly or not. Supplementary Figure 1e-f shows that an oscillating cell is able to repolarise completely during reversals and hence to migrate very persistently between the two ends of the footprint.

##### B. Oscillation analysis

Despite the wide variability in cellular behaviours, conserved patterns seem to emerge from the different cell trajectories. Therefore we resorted to a semi-manual analysis of the oscillations. Oscillation peaks were first automatically detected on the trajectory of the centre-of-mass, using an algorithm based on the Mexican hat wavelet transform. Then the output was manually corrected so as to remove false positive (mostly small amplitude undulations in zones with little movement). For each peak, the corresponding proximal extremum of the cell edge position was then searched, so that the final peak position corresponded to the actual extremal point reached by the cell edge, irrespective of the cell length that could vary a lot especially during reversals. Then we analysed the time series of the amplitude and period of the so-called "hemi-periods" (half of an oscillation, corresponding to one peak-to-peak excursion – the time period is doubled to correspond to the usual definition of a full oscillation). We noticed that most of the time a robust linear relation $A \propto T$ emerges, pointing to constant speed motion despite the long-term variation of oscillation amplitude. The time series of amplitudes were also consistently increasing with time, suggesting the existence of an extending accessible domain. Moreover, despite consistent change of slope at $A \simeq 70 - 100\,\mu\mathrm{m}$, the $(A_i)$ sequences grew roughly linearly, which is consistent with the idea of the addition of a constant "discovered" area at each iteration. Finally, it is interesting to note that in spite of differences in the full time cell trajectories, the oscillations could share their properties locally. It is well examplified in Supplementary Figure 2 where $A(T)$ and $(A_i)$ plots locally super-impose when the oscillations are in the same dynamical range – $t \in [30; 50]$ h for cell 1, $t \in [10; 30]$ h for cell 2 – while the two cells' history before and after differ completely.

##### C. The phase-space of linear cell migration.

Here we sought to extend the approach taken in the work by Bruckner *et al.* [5] for a case in which space is not bounded but footprint effects are taken into account. The idea is to draw a generic description of the cell behaviour by analysing the motion through the measurement of average and standard deviation of acceleration at any point in a proper phase space. The main task in that context is to replace the space variable – in unbounded space, $x$-invariance should be warranted – by suitable footprint variables. Both the speed and $\Delta\varphi$, the difference in footprint value between the two cell ends, showed significant correlations with the cell acceleration. To get a better estimate of the relation between $\Delta\varphi$ and $a$, we represented $a$ as function of both $\Delta\varphi$ and $\varphi_c$, the value at the cell centre. It appeared that $\langle a\rangle$ took significant values mostly at $\|\Delta\varphi\| = \varphi_c$, with an acceleration oriented towards higher $\varphi$ values (*ie* towards the interior of the footprint). This is shown in Supplementary Figure 4-5, with smoothed heatmaps highlighting the nonzero values of $\langle a\rangle$. This suggested that the footprint had an effect on the cell motion primarily near the footprint edges ($\|\Delta\varphi\| \simeq \varphi_c$ implies that $\varphi \simeq 0$ on one of the cell ends). To investigate this further, we decided to represent $\langle a\rangle$ as a function of $v$, $\varphi_l$ and $\varphi_r$ (Supplementary Figure 6). This made clear that the main effect of the footprint was restricted to the small zones where $\varphi_l \leqslant \varphi_0$ or $\varphi_r \leqslant \varphi_0$, with a critical threshold value $\varphi_0 \simeq 5\,\mathrm{h}$. Following this observation, we distinguished between the interior of the footprint and its borders based on this criterium on $\varphi_{l,r}$ and studied further the relationship between $a$ and $v$ in the two zones. Within the footprint, the data seem to follow a linear relationship, $\langle a\rangle = v/\tau_{va}$, which is common in persistent random cell migration. On the edge of the footprint, $\langle a\rangle$ is oriented inwards irrespective of the sign of $v$ with a parabolic relation between the two variables.

To verify that the $a - \varphi$ relationship is not an artifact of the overall increase of $\varphi$ with time, we reproduced the same measurements on limited time windows. We found that the output does not depend on the time of measurement, confirming that the observed effects are real effects of $v$ and $\varphi$. Finally, the time traces of $v$, $\varphi_l$ and $\varphi_r$ in Supplementary Figure 7 illustrate the motion of two stereotypical cells in this phase space.

##### D. Analysis of experimental trajectories through the PSAW model.

We discretised the trajectories to focus on the statistics of motion reversals independently from the detailed dynamics of cell speed and shape fluctuations. The main issue was to determine the right space and time scales to define the discretisation grids. In the framework of the PSAW model, the most natural spatial scale is the cell size, but as seen in Supplementary Figure 1, it exhibits both slow growth and fast fluctuations. As a simpler proxy, we used the minimum of the cell size in time as a reference length, $L_{\mathrm{ref}} = \min_t L$, to define the spatial grid (Supplementary Figure 8b). With that convention and the time resolution of our experiments $\Delta t = 6\,\mathrm{min}$, at each time frame the cells either stay on the same site or jump to a neighbouring site. We redefined the time grid by considering only the jump events (Supplementary Figure 8c), ignoring that jump times vary, as shown by their distribution (Supplementary Figure 9d), with an average of $\langle t_j\rangle = 0.6\,\mathrm{h}$. As a self-consistency check, we simulated PSAWs in the range of parameters found experimentally, then we expanded time by randomly picking jump times in the experimental $t_j$ distribution and smoothed the trajectories in space using a sliding average over $2\,\mathrm{h}$. Then we applied the discretisation procedure to those artificial trajectories and checked that the output $(k, \beta)$ couples are consistent with the input ones. This procedure is also robust to the choice of $L_{\mathrm{ref}}$ as shown in Supplementary Figure 9f by the very slight variation of the ouptut $k$ distribution when the median of $L(t)$ or a single $L_{\mathrm{ref}}$ value for all cells was used instead of the minimum.

We found that the experimental $(k, \beta)$ points are concentrated around the $\beta = -2(k + 1)$ line irrespective of the track width. Since it means that the reversal probability on the footprint edge is constant, it suggests that the main difference between oscillating and static cells resides in their ability to polarise persistently rather than in the self-attraction strength of their trajectory. Interestingly, the $k$ distribution is slightly shifted for $W = 10\,\mu\mathrm{m}$ which also displays less oscillating cells compared to the larger tracks. On conditioned substrates on the other hand, the $(k, \beta)$ scatter is moved to higher $\beta$ values, with a non-negligible fraction of positive $\beta$, which evidences a reduced effect of the cell path's self-attraction in that condition.

#### III. Supplementary Model

This Supplementary Material presents definitions of the random walk models discussed in the main text, and summarizes their main properties. We discuss

- the class of attractive self interacting random walks
- the SATW and PSATW processes for $d = 1$
- the SATW process for $d > 1$

##### A. Attractive self interacting random walks

In this section we give the definitions of the self interacting random walks of the attractive class discussed in the main text, and summarize their important properties.

##### 1. definitions

Self interacting random walks can be defined as nearest neighbor random walks on a $d$-dimensional lattice, for which the probability to jump to a neighboring site $i$ at time $t$ is proportional to a weight function $w(n_i)$ that depends on the number of previous visits $n_i$ of the random walker to site $i$ up to time $t$. Note that alternatively the variable $n$ can be defined on bonds instead of sites ; this prescription will be called bond convention below, and will not be discussed in details. Denoting by $x_t$ the position of the random walker at time $t$ and by $V(x_t)$ the set of neighboring sites, the transition probability defining the process can be written for $k \in V(x_t)$:

$$p(x_{t+1} = k|\{x_t, ..., x_0\}) = \frac{w(n_k)}{\sum_{j\in V(x_t)} w(n_j)}. \tag{1}$$

Note that the set of local times (or number of visits) $\{n_i\}_{i\in\mathbb{Z}^d}$ at time $t$ depends on the full trajectory $\{x_t, ..., x_0\}$ up to time t. The process is therefore strongly non Makovian and has long range memory. We focus in this paper on the attractive case, which is realized by weight functions $w$ that are monotonically increasing with $n$. Previous works have investigated exponential $[w(n) = e^{-\beta n}, \beta < 0]$, subexponential $[w(n) = e^{-\beta n^k}, \beta < 0]$, polynomial $[w(n) = n^\beta, \beta > 0]$ or asymptotically constant $[w(n) \sim 1]$ weight functions [ [11–15, 19, 21]]; some of the properties of these processes are reminded below.

##### 2. self-trapping

Qualitatively, attractive self interacting random walks are attracted by their own path. Strikingly, this can lead to the full trapping of the walker within a finite set of sites in the $t \to \infty$ limit, and therefore to a bounded mean squared displacement (MSD). This effect was demonstrated mathematically by B. Davis [11] for 1–dimensional attractive self interacting random walks with the bond convention. This result states that if $\sum_{n=1}^{\infty} w(n)^{-1} = \infty$ the random walker is free and will visit infinitely often all the sites of the lattice. Conversely, for $\sum_{n=1}^{\infty} w(n)^{-1} < \infty$, the random walker visits only a finite set of sites and will eventually almost surely oscillate between two adjacent sites. Even if no mathematical proof is available, these results are conjectured to apply more generally to the site convention that we use in this paper, and for $d$–dimensional lattices [19]. Finally, this indicates that exponential or subexponantial or polynomial (with $\beta > 1$) attractive self interacting random walks lead to the full trapping of the random walker and to a bounded MSD. We focus below on the class of free attractive self interacting random walks, for which the MSD diverges for $t \to \infty$.

##### 3. free attractive self interacting random walks

In this section we provide a brief survey of attractive self interacting random walks that are free (ie have diverging MSD for $t \to \infty$), before focusing on the models that are used in paper (SATW and PSATW).

*a. Polynomial self interacting random walks*

This model is defined by the transition probability (1) with the weight function [20, 21]:

$$w(n) = \frac{1}{|1-\alpha|}(n/2)^\alpha - \frac{B}{(1-\alpha)^2}(n/2)^{\alpha-1} + O(n^{\alpha-2}) \text{ where } B \in \mathbb{R}. \tag{2}$$

So far it has been studied mostly for $d = 1$. The Davis theorem stated above and its generalization show that for $1 < \alpha$ the random walker is fully trapped. For $0 < \alpha < 1$ the walk is free (the MSD diverges for $t \to \infty$), and the scaling of the MSD has been determined in $1d$ for the bond convention : $\langle x_t^2 \rangle \sim t^{\frac{2(1-\alpha)}{2-\alpha}}$. Note that numerical results indicate that this scaling does not hold for the site convention. The persistence exponent is not known for this process, as well as the scaling of the MSD for $d > 1$.

*b. Self interacting random walks with bounded weights : the self attracting walk (SATW)*

This model is defined by the transition probability (1) with the weight function [9, 10, 16, 18]:

$$w(n) = \exp(-\beta f(n)), \tag{3}$$

where $f(0) = 0$ and $f(n > 0) = 1$ and $\beta < 0$. In this regime, for $d = 1$, the random walk is free and the scaling of the MSD is diffusive [17] : $\langle x_t^2 \rangle \sim t$. This model, with its generalization to persistent random walks discussed below, is the central model on which we focus in this paper. This choice more generally covers all cases where $f$ is bounded. For practical applications, and in particular in the context of cell migration that we study in this paper, it means that either the deposited signal saturates, or the cell response to the deposited signal saturates. While such hypothesis –even if realistic– cannot be directly challenged experimentally, our experimental results are consistent with this model. Of note, the SATW dynamics at time $t$ is fully determined by the position of the walker $x_t$ and the visited territory $\mathcal{D}_t$, ie the set of sites that have been visited up to time $t$. For $d=1$, $\mathcal{D}_t$ is a connected segment, which greatly simplifies the analysis as compared to $d>1$. When the walker is inside $\mathcal{D}_t$ (away from its boundaries) the dynamics is that of a symmetric Polya walk ; the dynamics is modified only when the walker reaches the boundary of $\mathcal{D}_t$. The properties of the SATW model will be discussed in more details in the next sections.

*c. The persistent self attracting walk (PSATW)*

To take into account the persistence of migrating cells, we make use in this paper of a generalization of the SATW called persistent self attracting walk (PSATW) [6]. For $d=1$, it is defined as follows. When the walker is on a site $i$ within the visited domain $\mathcal{D}_t$ – ie surrounded by sites that have already been visited, $n_{i-1}, n_{i+1} > 0$ – it performs a classical persistent random random walk : it changes direction with probability $p_{r,i} = \frac{e^{-k}}{e^{-k}+e^{k}}$, and reproduces its previous step with probability $1-p_{r,i}$. Here $k>0$ is a parameter that controls the cell persistence length $l_p = e^{2k}$, or equivalently persistence time $t_p = l_p$ (the speed is set to 1 in this discrete model). When the walker is at an edge of the domain $\mathcal{D}_t$ (eg $n_{i+1}>0$), it experiences a local bias inward the visited domain parametrized by $\beta<0$ and the probability to change direction can be written $p_{r,e} = \frac{e^{-k-\beta}}{e^{-k-\beta}+e^{k}}$, while the probability to reproduce the previous step is $1-p_{r,e}$. In particular for $k=0$ the SATW model is recovered ; for $\beta=0$ the classical persistent random walk is recovered. For $d=1$, the random walk is free and the scaling of the MSD is diffusive in the long time limit [6] : $\langle x_t^2\rangle \sim t$. The properties of the PSATW model will be discussed in more details in the next sections.

*d. Asymptotically free walk :*

This model is defined by the transition probability (1) with the weight function [20, 21] :

$$w(n) = 1 - 2Bn^{-1} + O(n^{-2}) \text{ where } B \in \mathbb{R}. \tag{4}$$

It was introduced as a refinement of the SATW ; for $d=1$, the MSD is diffusive as for the SATW. The persistence exponent is not known for this process, as well as its properties for $d>1$.

##### B. SATW and PSATW for $d=1$.

In this section we review in more details the properties of the SATW and PSATW models for $d=1$, which are used in the main text (see Fig 12).

###### 1. MSD and ageing of increments

The MSD of both SATW and PSATW models for $d=1$ are diffusive [6, 17] for $t\to\infty$. We discuss in this section the ageing properties of the increments $\langle [x(t+T)-x(T)]^2\rangle$.

*a. SATW*

The increments of the SATW have been shown to display scale free ageing [6]. It means that in the limit $t, T \gg 1$ the increments can be written

$$\langle [x(t+T)-x(T)]^2\rangle = 2D(t/T)\,t, \tag{5}$$

where the diffusion coefficient $D(t/T)$ has the following finite limits :

$$\begin{cases} D(t/T) \sim D_L(\beta) \text{ for } t \gg T \\ D(t/T) \sim D_s \text{ for } t \ll T. \end{cases} \tag{6}$$

The long time diffusion coefficient $D_L(\beta)$ is not known analytically ; however, $t \ll T$ it is easy to find that $D_s = 1/2$, because the random walker spends most of its time inside the visited domain $\mathcal{D}_t$, and thus performs a symmetric Pólya walk. Ageing of the increments is clearly seen by the explicit dependence of the increments on the observation time $T$ (see Fig 12).

*b. PSATW*

The increments of the PSATW can be deduced from the above analysis of the SATW [6]. In the long time limit $t \gg T, t_p$ the scaling of the increments is not modified by persistence :

$$\langle [x(t+T)-x(T)]^2\rangle \sim 2D_L^p(\beta)\,t. \tag{7}$$

In the regime $t \ll T$ one recovers the classical behavior of persistent random walks. For $t \ll t_p$ the scaling is ballistic

$$\langle [x(t+T)-x(T)]^2\rangle \sim\ t^2, \tag{8}$$

with a cross over to a diffusive regime for $t \gg t_p$

$$\langle [x(t+T)-x(T)]^2\rangle \sim 2D_s\,t, \tag{9}$$

with $D_s = e^{2k}/2$. Ageing of the increments is clearly seen by the explicit dependence of the increments on the observation time $T$ (see Fig 12).

##### 2. *First-passage properties : persistence.*

We conclude this section by reminding the first passage properties of the SATW and PSATW models for $d = 1$. We define the survival probability $S(t)$ as the probability that the walker has not reached a target at time $t$. The large time behaviour of the survival probability is characterized by a power law decay $S(t) \propto t^{-\theta}$ that defines the persistence exponent [8] $\theta$. The persistence exponent was shown [6] to be given for both the SATW and PSATW by

$$\theta = e^{-\beta}/2. \tag{10}$$

In particular one has $\theta > 1/2$. As compared to Brownian random walks ($\theta = 1/2$), the relative weight of short trajectories with respect to long trajectories is thus increased : exploration is thus more local.

#### C. SATW model for $d > 1$

In this section, we discuss the properties of the SATW, and more precisely the scaling of the MSD and of its increments; these properties are also expected to apply to the generalised PSATW model in the regime $t, T \gg t_p$. As for $d = 1$, the dynamics of the SATW model is fully defined by the position of the random walker $x_t$ and visited territory $\mathcal{D}_t$ at time $t$. Qualitatively, the random walker is attracted by $\mathcal{D}_t$: for $d > 1$, the geometry of $\mathcal{D}_t$ is however complex, and only few exact results are available for this process, which has been studied mostly numerically. To determine the scaling of the MSD, a key ingredient is the growth rate of $\mathcal{D}_t$ ; we provide below scaling arguments similar to [14] to derive these scalings for $d = 3$ and $d = 2$ ; of note the case $d = 2$ is still debated [13, 14] .

##### 1. *MSD for* $d = 3$

We start with the simplest case $d = 3$ (see Fig.14). We write

$$\frac{d\langle \mathcal{D}_t \rangle}{dt} \sim \frac{1}{\langle T \rangle_s} \tag{11}$$

where $\langle T \rangle_s$ is the mean return time to the boundary of $\mathcal{D}_t$ (we identify $\mathcal{D}_t$ and its volume), averaged over all starting positions on the boundary of $\mathcal{D}_t$. Making use of the so called Kac formula for mean return times [7], the scaling of $\langle T \rangle_s$ can be written

$$\frac{1}{\langle T \rangle_s} \sim \frac{\delta \mathcal{D}_t}{\mathcal{D}_t} \tag{12}$$

where $\delta \mathcal{D}_t$ denotes the boundary of $\mathcal{D}_t$. We now define the walk dimension $d_w$ of the process by the scaling of the MSD: $\langle x^2 \rangle \sim t^{2/d_w}$. We also introduce $d_{fc}$ and $\alpha$ as the fractal dimensions of $\mathcal{D}_t$ and of its boundary $\delta \mathcal{D}_t$ respectively. Using that the length scale of $\mathcal{D}_t$ and $\delta \mathcal{D}_t$ is $\sqrt{\langle x^2 \rangle}$, this yields :

$$\mathcal{D}_t \sim t^{d_{fc}/d_w}, \tag{13}$$

$$\delta \mathcal{D}_t \sim t^{\alpha/d_w}. \tag{14}$$

Using the scalings (13) and (14) in (11) and (12) leads to the following condition of self-consistency:

$$\alpha = 2d_{fc} - d_w. \tag{15}$$

We first assume that the strength of the self interaction is small enough ($|\beta|$ small) so that $\mathcal{D}_t$ has the same scaling properties as for a classical symmetric random walk for $d = 3$. It is known that the territory explored by a classical random walk is characterized by $\alpha = d_{fc} = 2$. Equation (15) is then satisfied for $d_w = 2$. This scaling argument suggests that for $|\beta|$ small, the SATW displays a normal diffusive scaling, which is indeed observed numerically. Let us now assume that for $|\beta|$ large the dimension of $\mathcal{D}_t$ is modified ; more precisely we assume that $\mathcal{D}_t$ is a smooth volume of dimensions $d_{fc} = d = 3$ and $\alpha = d_{fc} - 1 = 2$. Equation (15) then leads to $d_w = 4$. This scaling argument therefore suggests the existence of a sub-diffusive regime with $\langle x^2(t) \sim \rangle \sim t^{1/2}$. This subdiffusive regime is indeed observed numerically. Of note, the transition between the diffusive and subdiffusive regimes is discussed in [13].

##### 2. *MSD for* $d = 2$

In principle, the above scaling analysis can be reproduced for $d = 2$. For $|\beta|$ large, it suggests a subdiffusive regime $\langle x^2(t) \rangle \sim t^{2/3}$, which is indeed observed numerically. The behaviour for $|\beta| \to 0$, and in particular the transition to a diffusive regime is however still debated [13, 14] (see Fig.13) .

*3. Increments*

We discuss in this section the aging properties of the increments $\langle [x(t+T) - x(T)]^2 \rangle$.

*a. subdiffusive regime ($|\beta|$ large)*

The numerical analysis of the process suggests the following behaviours in the sub-diffusive regime for both $d = 2$ and $d = 3$ ($|\beta|$ large). For $t \gg T$ the scaling of the MSD is obtained : $\langle [x(t+T) - x(T)]^2 \rangle \sim t^{2/d_w}$, where we recall that $d_w = 4$ for $d = 3$ and $d_w = 3$ for $d = 2$. For $t \ll T$, the random walker is within $\mathcal{D}_t$, which is a smooth $d$–dimensional volume ; the process is thus a classical diffusion and $\langle [x(t+T) - x(T)]^2 \rangle \sim 2dD_s t$. Ageing of the increments is clearly seen by the explicit dependence of the increments on the observation time $T$ (see Fig 13,14).

*b. diffusive regime ($|\beta|$ small and d=3)*

Our numerical analysis suggests that the increments are asymptotically stationary for $d = 3$ in the diffusive regime (see Fig 14) :

$$\langle x^2(t) \rangle \sim 2D_L t. \tag{16}$$

*4. Impact on exploration : type problem and FPT properties*

The existence of a sub diffusive regime for $d = 2, 3$ has drastic consequences on space exploration. Indeed, in the subdiffusive regime one hase $d_w > d$ and exploration is called compact for $d = 2, 3$, as opposed to the classical symmetric Polya walk, which is non compact for $d = 3$ and only marginally compact for $d = 2$. For compact random walks, each site of the lattice is visited ultimately infinitely many times, and all sites are eventually visited with probability 1. Qualitatively, the random walker explores densely and exhaustively its neighborhood, while $\mathcal{D}_t$ grows smoothly with ”small” fluctuations. In particular the resulting survival probability is observed to decay faster than any power law, so that the persistence exponent is effectively infinite in this regime (see Fig 13,14).

#### IV. Supplementary videos

**Supplementary Video 1** Static MDCK cell on a 20 µm track. Phase contrast images. Scale bar 100 µm.

**Supplementary Video 2** Oscillating MDCK cell on a 20 µm track. Phase contrast images. Scale bar 100 µm.

**Supplementary Video 3** MDCK cell on a 20 µm track, alternating between oscillatory and static phases. Fluorescence intensity of PBD-YFP. Scale bar 50 µm.

**Supplementary Video 4** Oscillating MDCK cell on a 20 µm track. Fluorescence intensity of PBD-YFP. Scale bar 50 µm.

**Supplementary Video 5** MDCK cell on a control (top, same video as Supplementary Video 1) and a conditionned (bottom) 20 µm track. Scale bars 100 µm.

#### V. Supplementary Figures

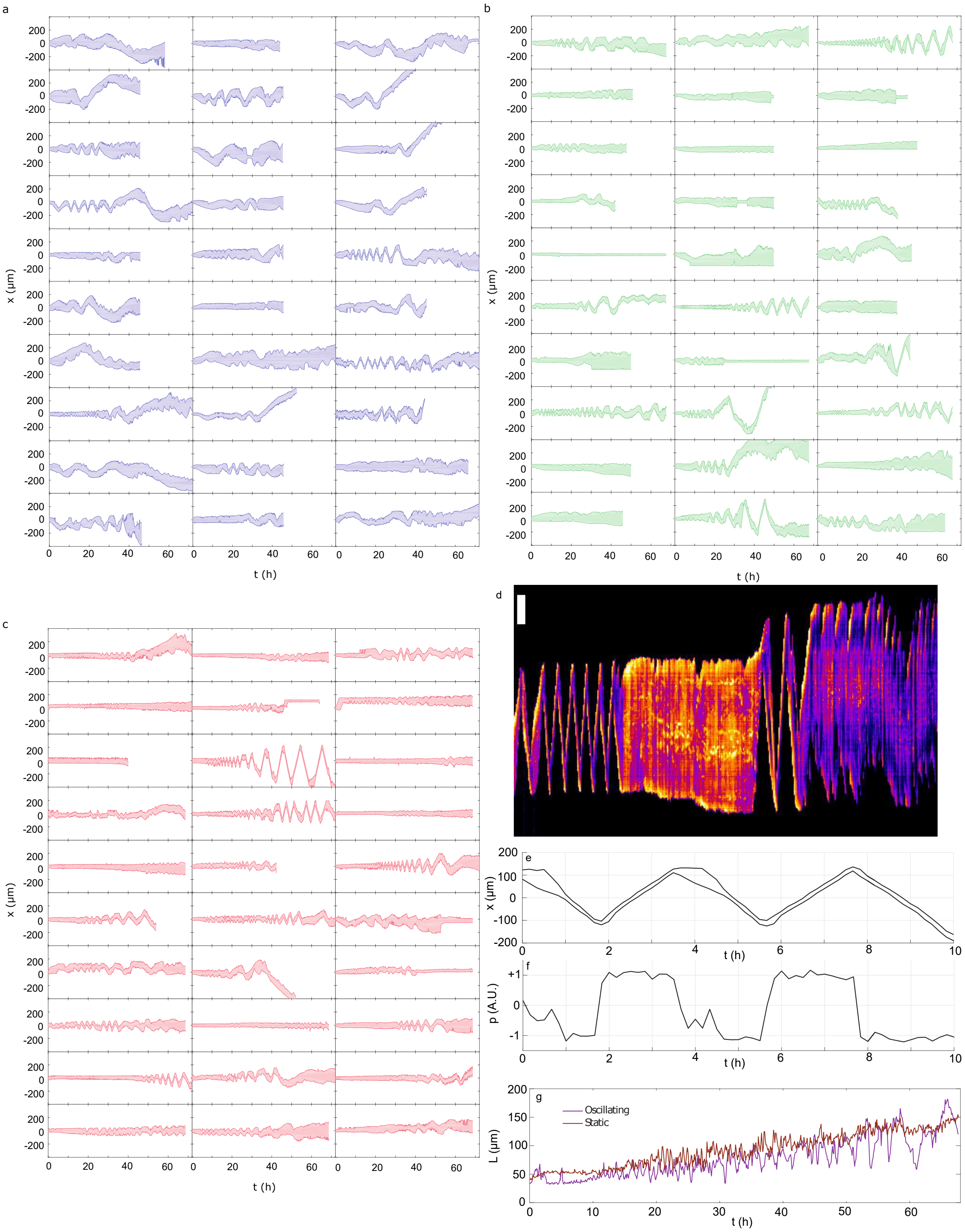


Supplementary Figure 1. **a–c.** 30 randomly sampled trajectories of MDCK cells on linear tracks of width $W = 10$ (**a**), 20 (**b**) and 50 $\mu$m (**c**) respectively. The vertical $x$ scale is 200 $\mu$m per tick, the horizontal $t$ scale is 10 h per tick (full ranges, $t \in [0; 72]$ h, $x \in [-400; 400]$ $\mu$m). **d.** Cell length as a function of time for the two examples showed in Figure 1d-e. **e.** Edge position as function of time for the PDB-YFP MDCK cell showed in Figure 1g. **f.** Polarity $p$ as a function of time for the PBD-YFP MDCK cell showed in Figure 1g. $p$ is computed as the moment of the spatial distribution of PBD signal intensity in the cell, with respect to the cell centre-of-mass. **g.** Kymograph of the PBD signal in a MDCK cell undergoing oscillations and static spreading in alternation. Scale bar 20 $\mu$m

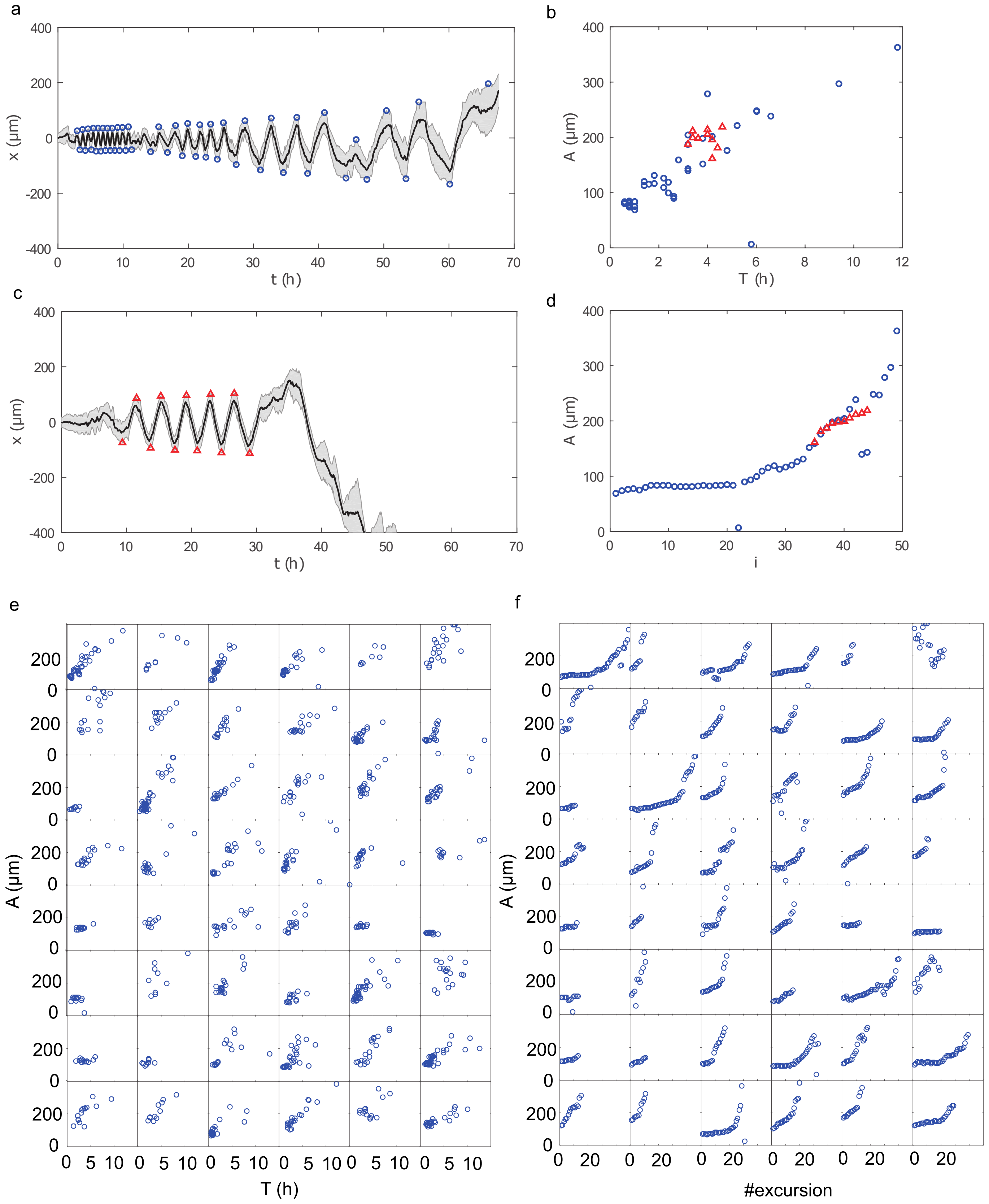


Supplementary Figure 2. Oscillation analysis. **a-b.** Two cell kymographs with the detected peaks marked as blue circles (**a**) and red triangles (**b**). **c.** Amplitude versus period of detected oscillations for the two examples in a (blue circles) and b (red triangles). d. Sequence of amplitude of oscillations in the two examples in a and b. The data from panel b have been shifted to the right to show that their rate of increase is similar to that of oscillations of same amplitude in a. **e.** Amplitude of oscillations as a function of their time period. Each plot corresponds to an individual trajectory. $W = 20\ \mu\mathrm{m}$. **f.** Time series of the amplitude of oscillations. Same data as in **e**.

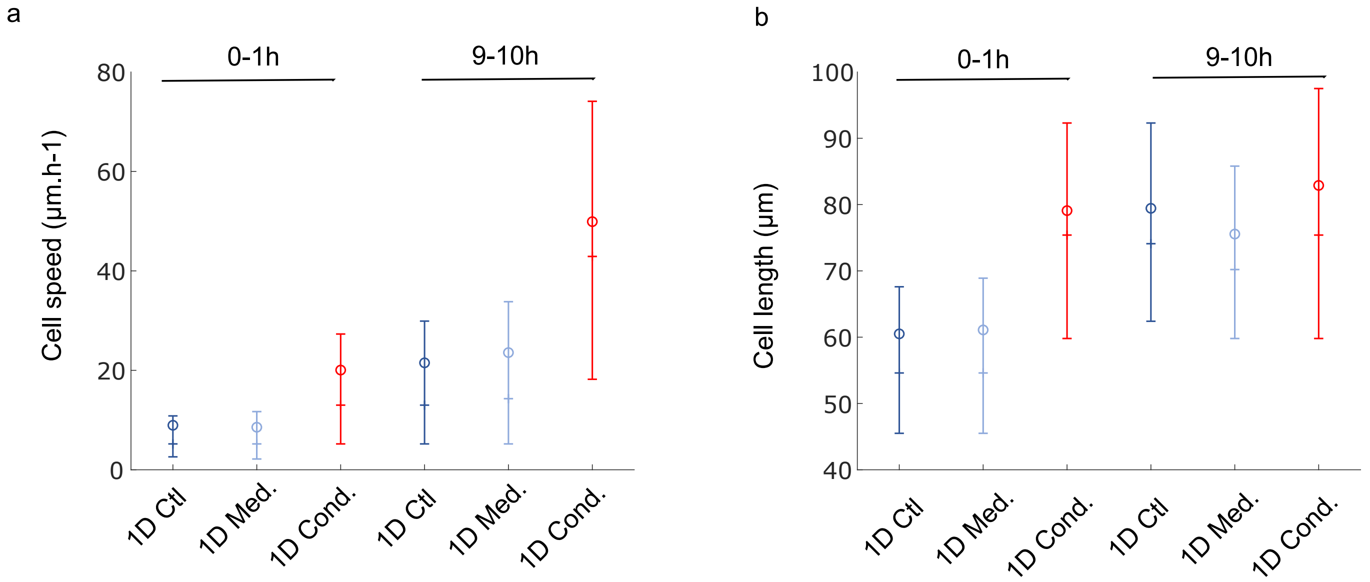


Supplementary Figure 3. Effect of conditioning on cell characteristics. **a.** Instantaneous cell speed for MDCK cells on control, control with medium and conditioned linear substrates, at early ($0-1$ h) and later ($9-10$ h) times. **b.** Cell length for MDCK cells on control, control with medium and conditioned linear substrates, at early ($0-1$ h) and later ($9-10$ h) times.

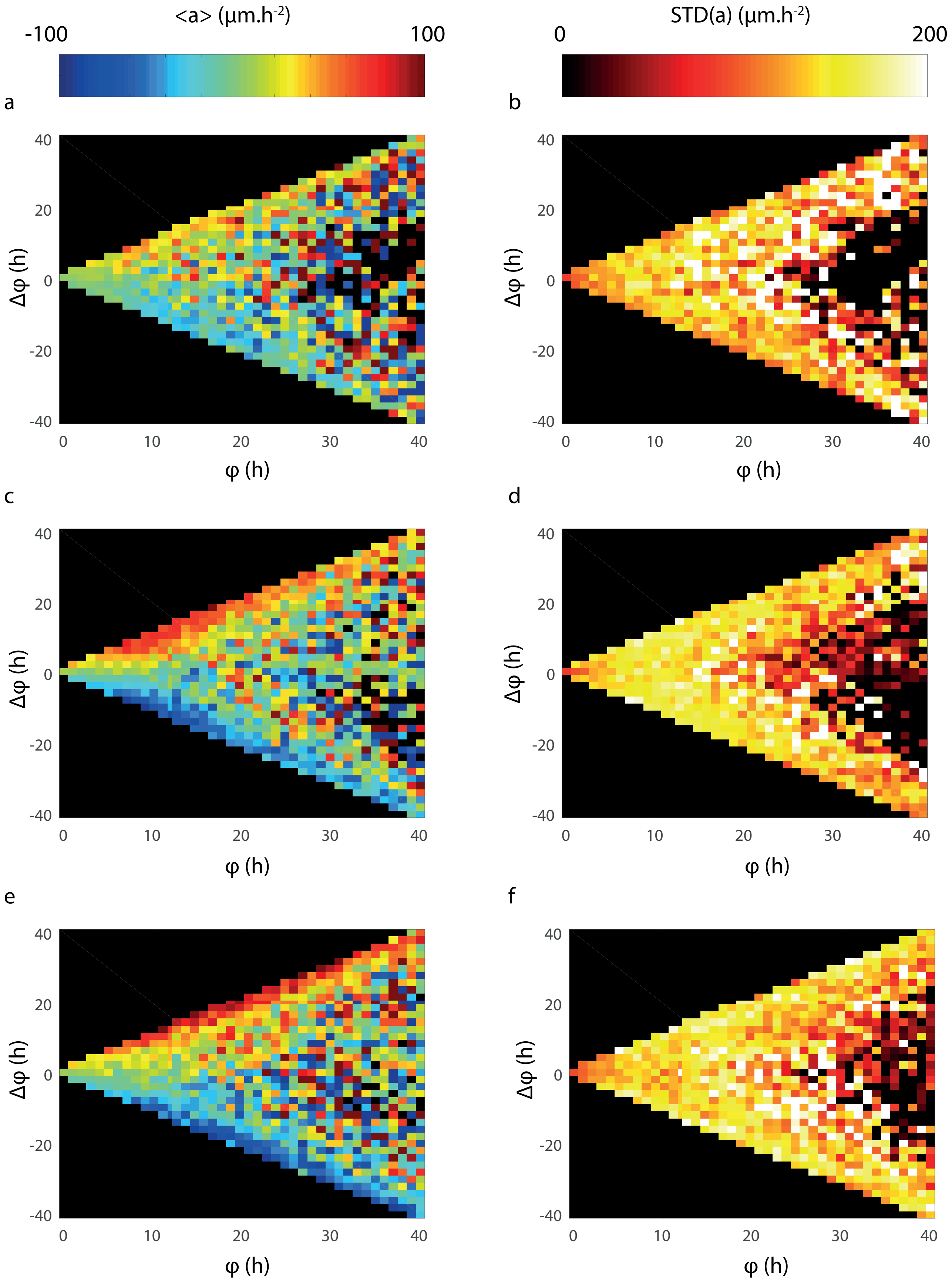


Supplementary Figure 4. $\varphi - \Delta\varphi$ phase space. Average (**a, c, e**) and standard deviation (**b, d, f**) of the acceleration as a function of $\varphi$ and $\Delta\varphi$ for cells on tracks of width 10 (**a,b**), 20 (**c,d**) and 50 (**e,f**) μm.

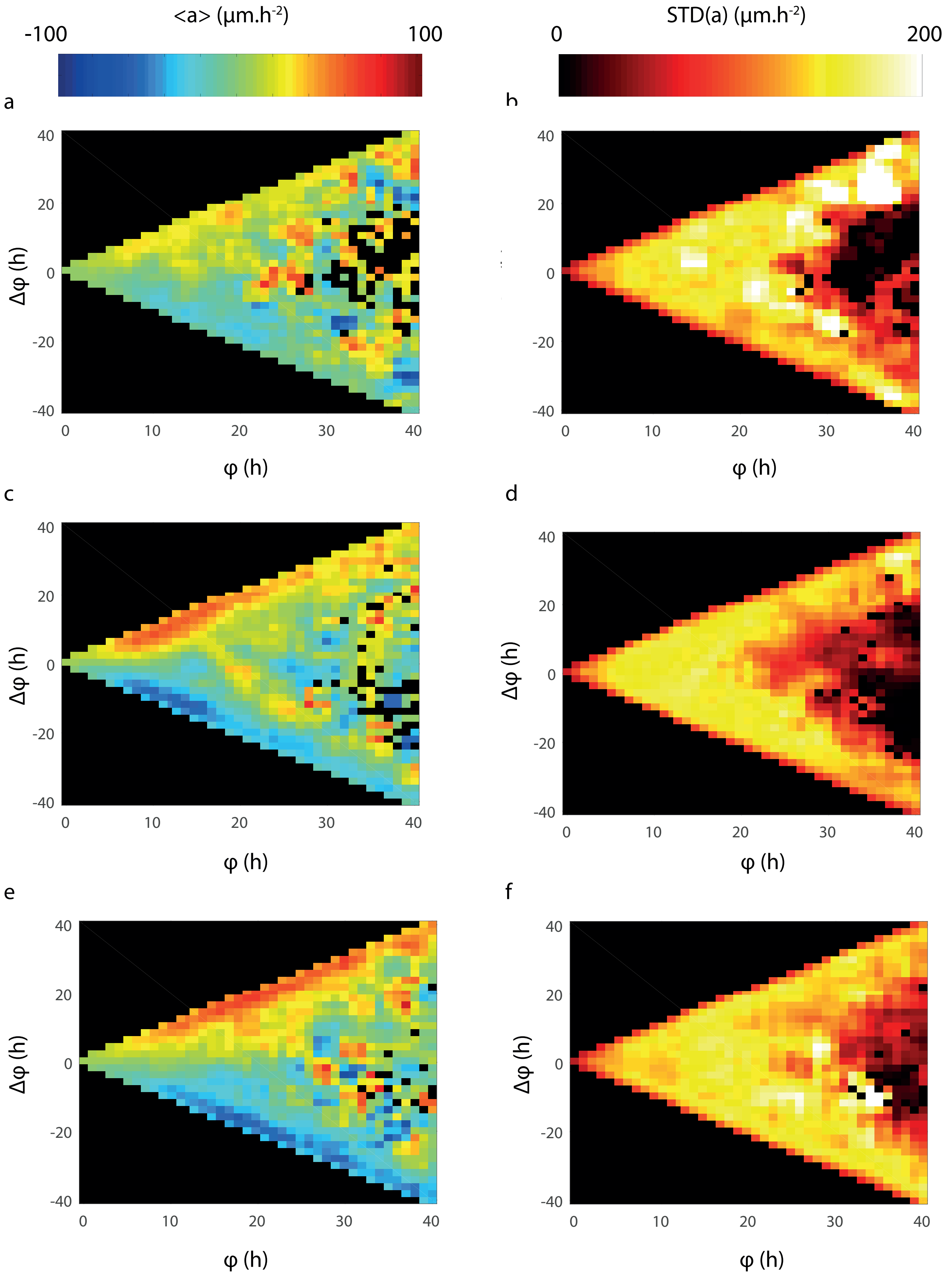


Supplementary Figure 5. $\varphi - \Delta\varphi$ phase space. Average (**a, c, e**) and standard deviation (**b, d, f**) of the acceleration as a function of $\varphi$ and $\Delta\varphi$ for cells on tracks of width 10 (**a,b**), 20 (**c,d**) and 50 (**e,f**) $\mu$m. Same data as in Supplementary Figure 4, smoothed over 3x3 squares.

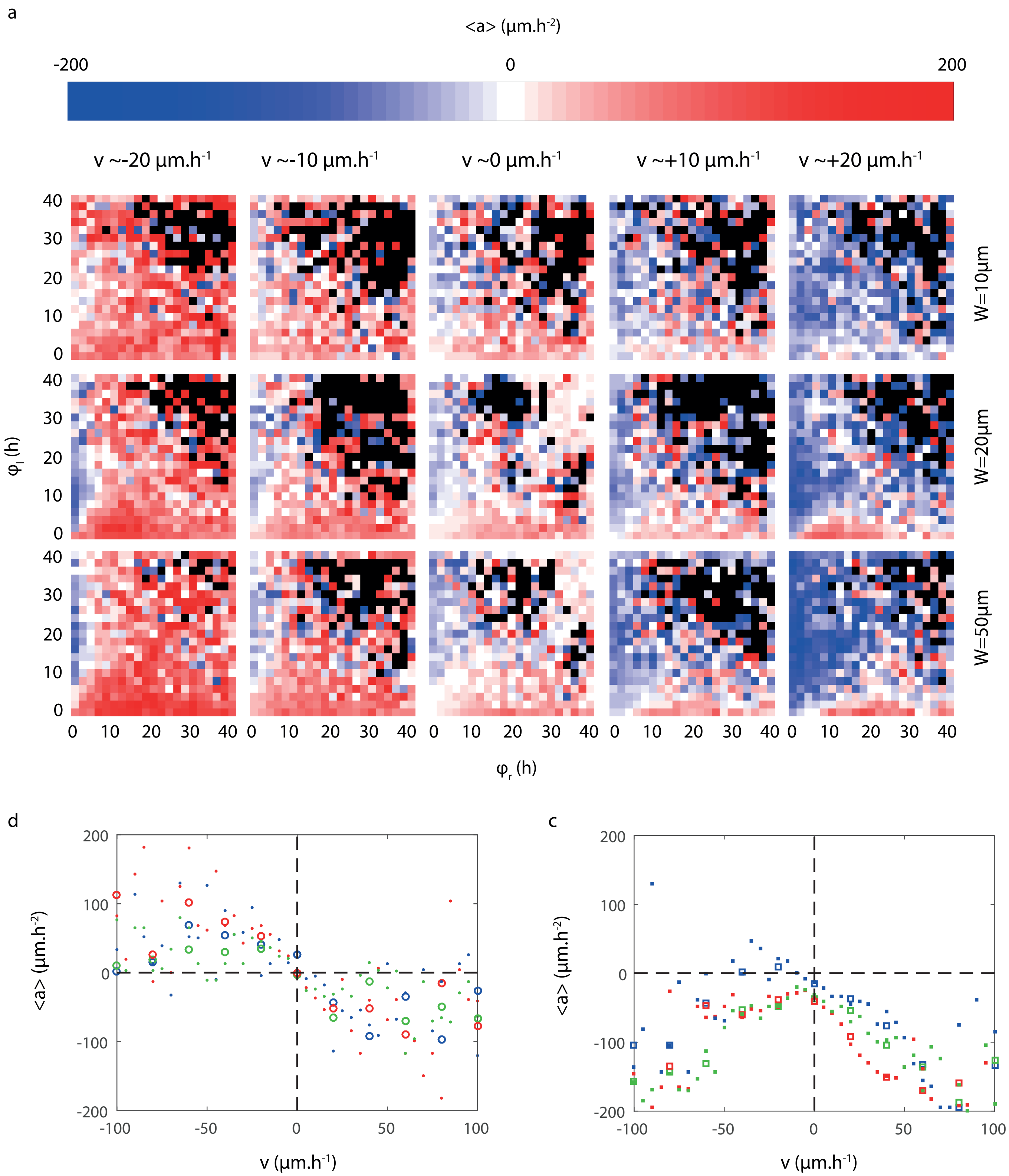


Supplementary Figure 6. $\varphi_l - \varphi_r$ phase space. **a.** Average acceleration as a function of $v$, $\varphi_l$ and $\varphi_r$ for cells on tracks of various widths. **b.** $\langle a \rangle$ as a function of $v$ measured near the $\varphi_l = \varphi_r$ diagonal. **c.** $\langle a \rangle$ as a function of $v$ measured near the $\varphi_r = 0$ axis (data near the $\varphi_l = 0$ axis also pooled after symmetrisation).

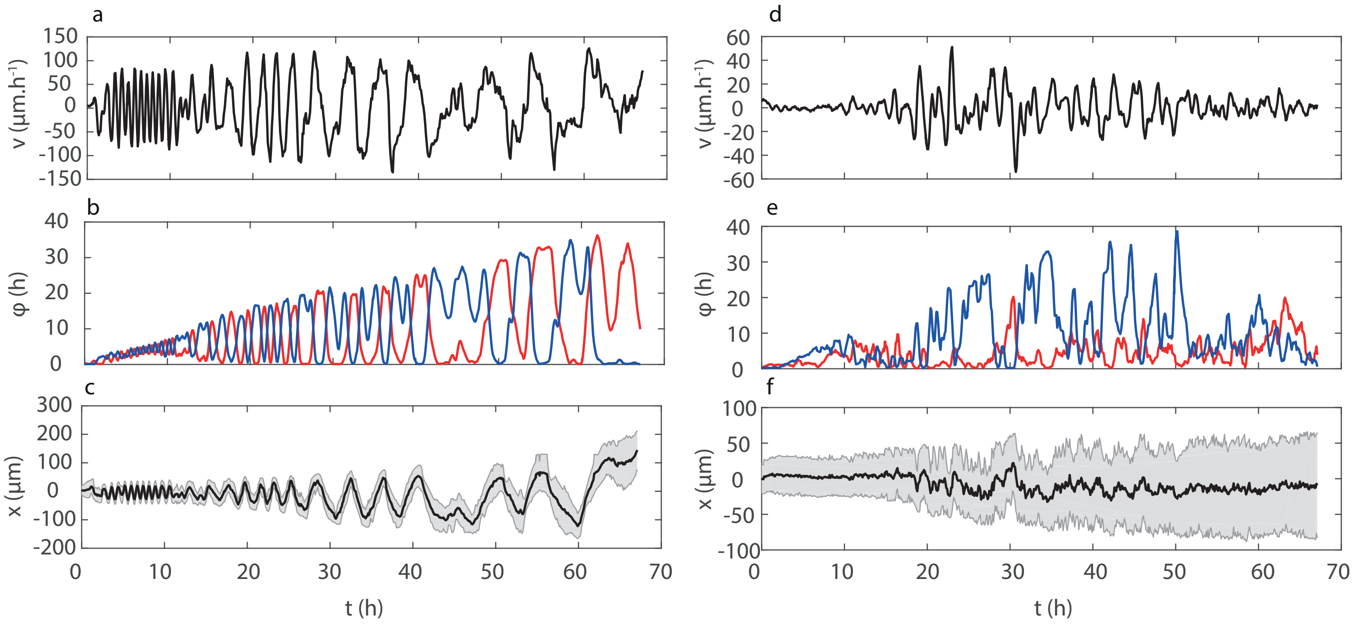


Supplementary Figure 7. Motion in the $(v, \varphi_l, \varphi_r)$ phase space. Velocity (**a,d**), footprint values at the cell ends (**b,e**) and kymographs (**c,f**) for the two cells shown in Figure 1d-e of the main text.

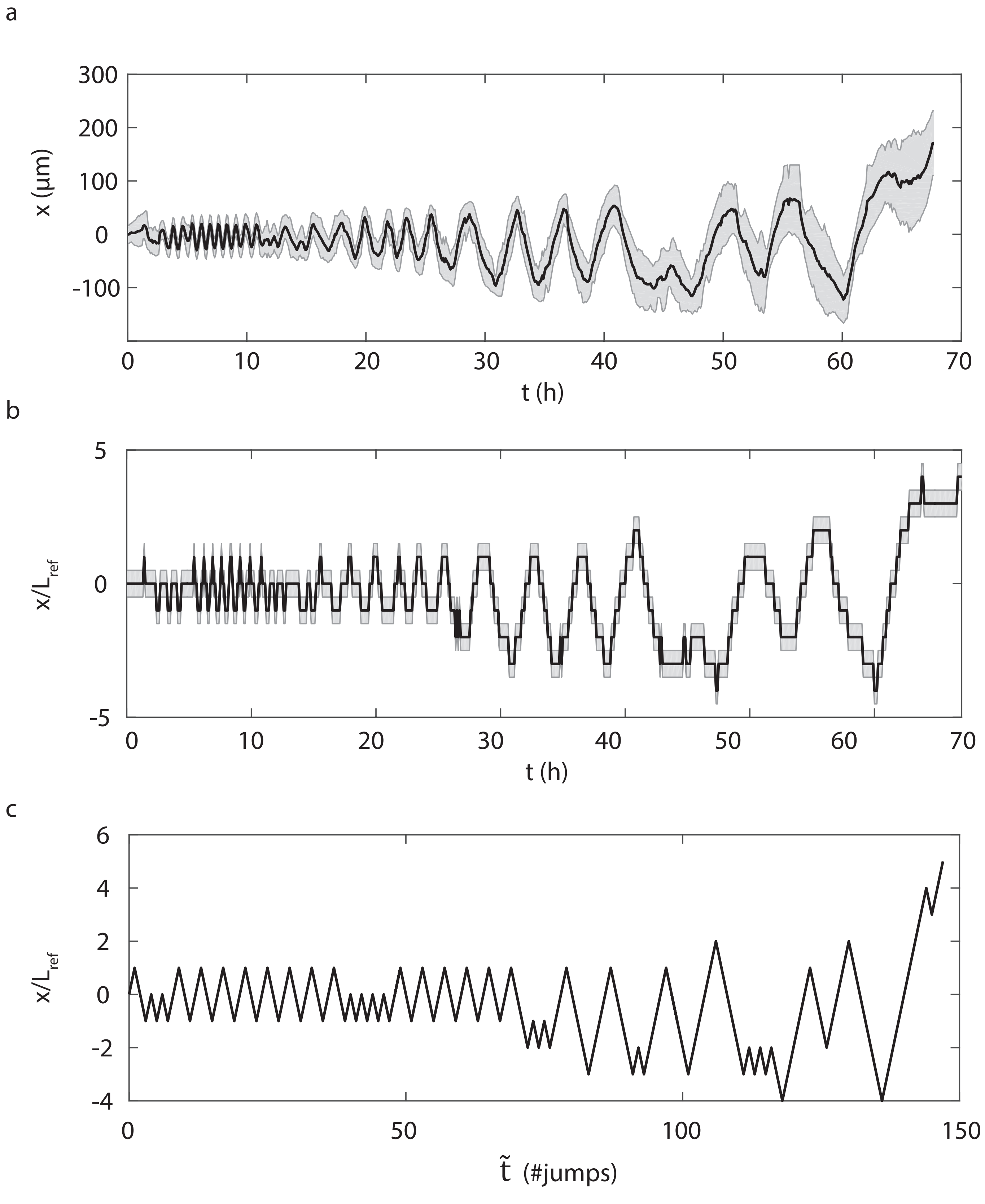


Supplementary Figure 8. Discretisation procedure. **a.** Original cell kymograph. **b.** Cell kymograph after discretisation of space using the cell's minimal length as $K_{\mathrm{ref}}$. **c.** Cell kymograph after keeping only jump events as time steps.

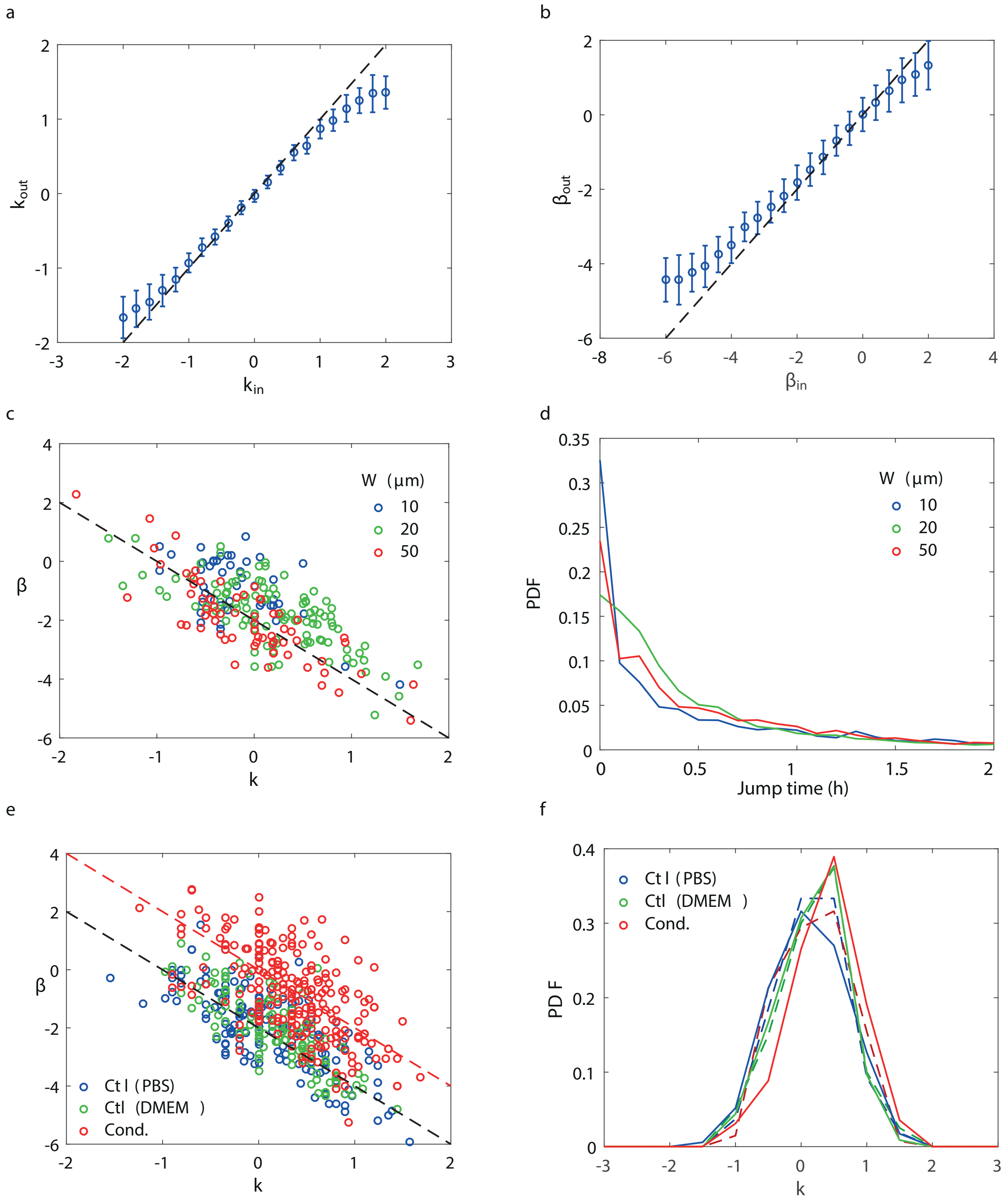


Supplementary Figure 9. PSAW analysis. **a.** Measured $k$ values as a function of input $k$ in simulated trajectories analysed with the discretisation procedure. **b.** Measure $\beta$ values as a function of input $\beta$ values in simulated trajectories analysed with the discretisation procedure. **c.** Experimental $\beta$ versus $k$ from cells on tracks of different widths. $\beta = -2(k+1)$ fit (dashed line). **d.** Distribution of jump times measured during the discretisation procedure for cells on tracks of different widths. **e.** Expermental $\beta$ versus $k$ for cells on control or conditioned substrates on lines of 20 $\mu$m in width. $\beta = -2(k+1)$ (black) and $\beta = -2k$ (red) fits (dashed lines). **f.** Distribution of measured $k$ for cells on control or conditioned substrates, with two definitions of $L_{\mathrm{ref}}$: minimal (solid lines) or median (dashed lines) of the cell length.

a
b

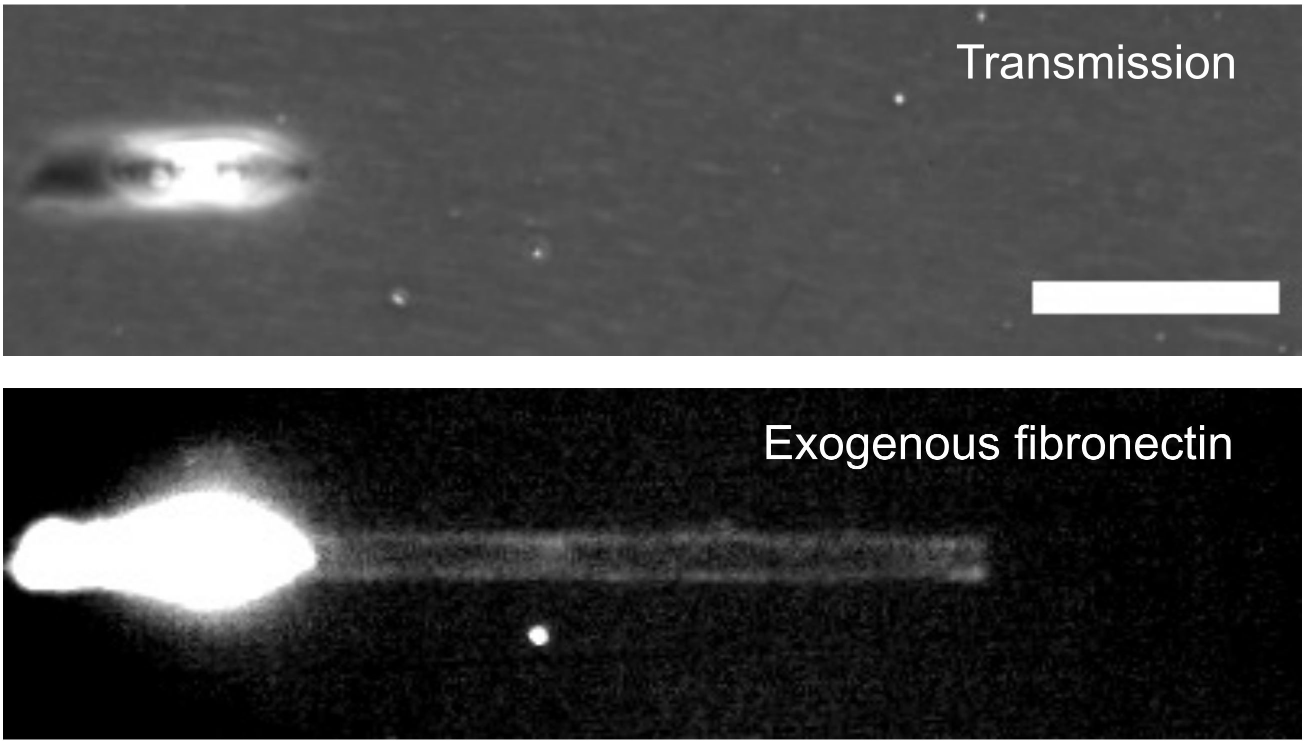


Supplementary Figure 10. Exogeneous fibronectin is captured and deposited by moving cells. **a.** Phase contrast image of an isolated cell on a 20 μm line after 48 h incubation with labeled fibronectin and fixation. **b.** Corresponding fluorescence image of the labeled fibronectin, showing signal along the line close to, but away from the cell. Scale bar 100 μm.

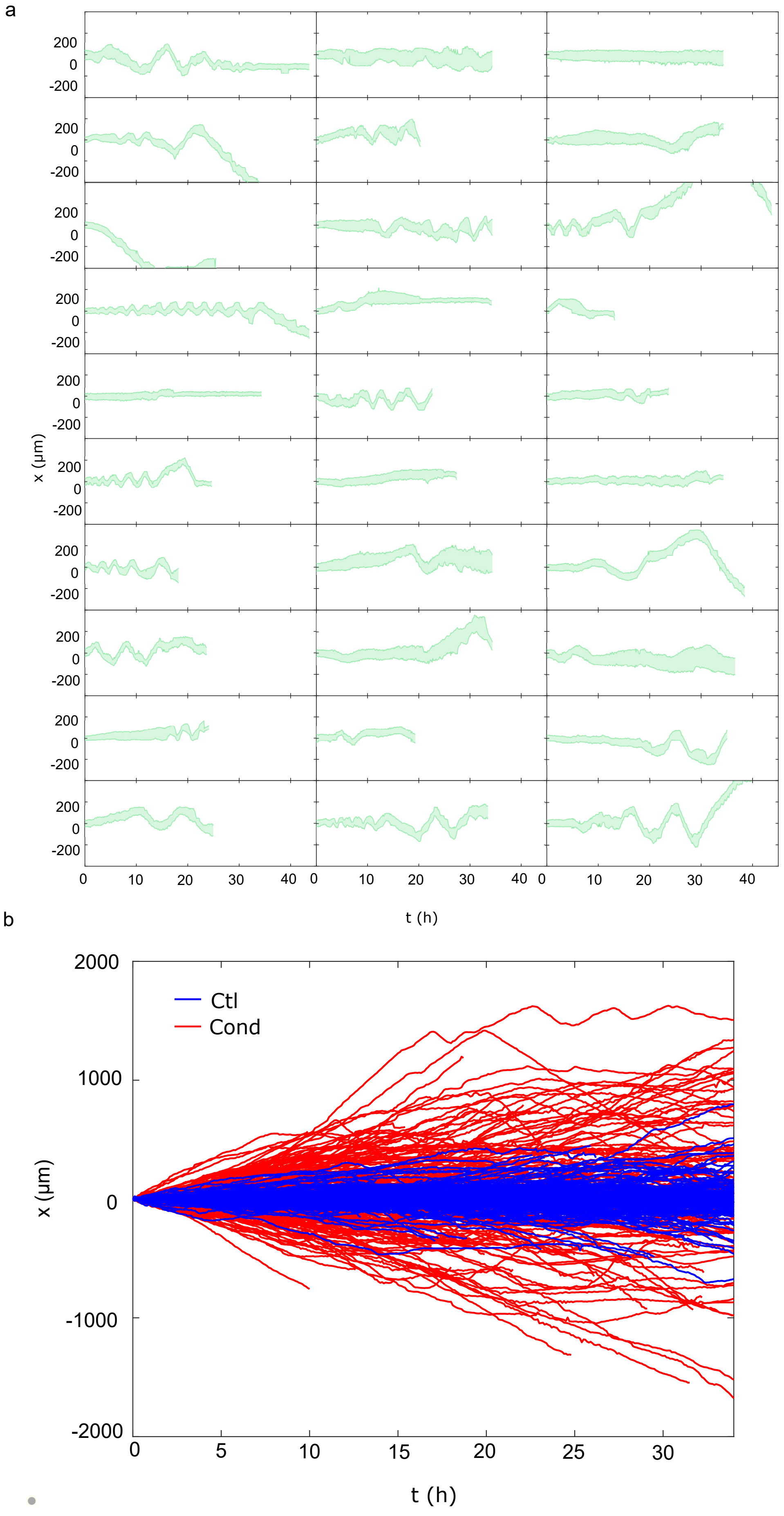


Supplementary Figure 11. Caco2 cells also exhibit footprint-related oscillations. **a.** Sample kymographs of isolated Caco2 cells on 20 µm line patterns. **b.** Trajectories of Caco2 cells on control (Ctl, blue) and conditioned (Cond., red) 1D substrates. Resp. $n = 246$ and 216 trajectories for Ctl and Cond. substrates from 3 independent experiments.

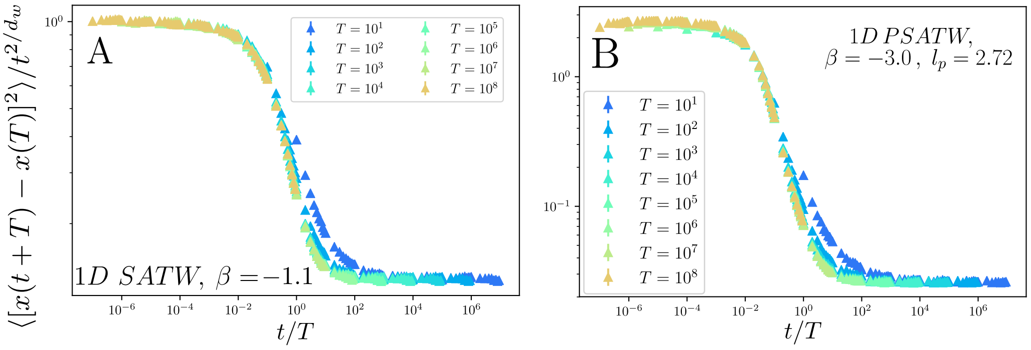


Supplementary Figure 12. a) b) Aging of the increments for the SATW and the PSATW (normalized by the expected diffusive scaling at long times). Each curve corresponds to a fixed value of $T$.

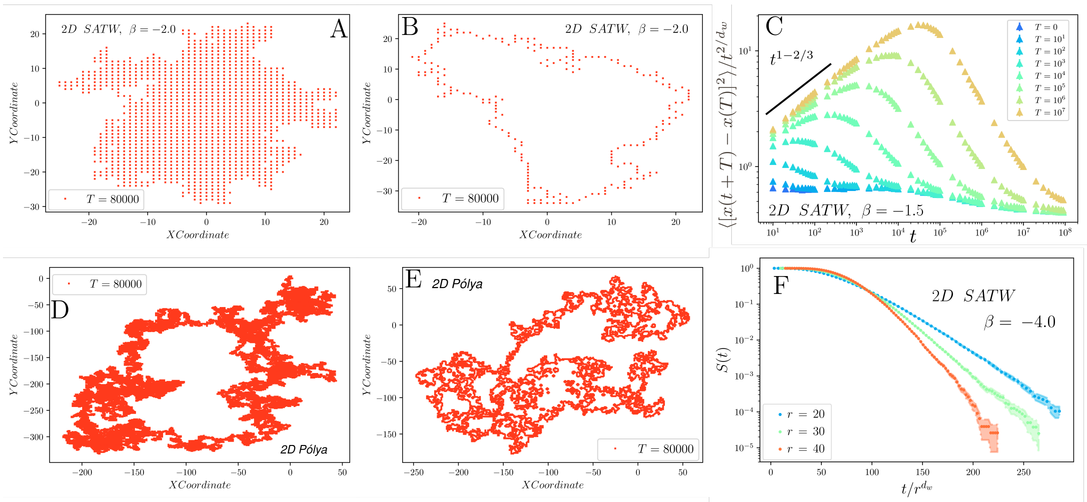


;

Supplementary Figure 13. Area visited and aging for the SATW in dimension 2, and comparison with a simple random walk. a) d) Area visited in dimension 2 for a subdiffusive SATW and for a simple random walk. The set of visited sites $\mathcal{D}_t$ for the SATW grows as a 2–dimensional smooth compact set with an area that scales as $t^{4/d_w}$ contrary to the simple random walk where $\mathcal{D}_t$ as a null area in the continuum limit. b) d) Boundary of $\mathcal{D}_t$ for the SATW random walk and for a simple random walk. The dimensions of the boundary sets are $\alpha_p = 1$ for the SATW and $\alpha_p = 2$ for a simple random walk. c) Aging of the increments for the subdiffusive SATW normalized by the expected diffusive scaling at long times. Each curve corresponds to a fixed value of $T$. NOte that the increments are diffusive for $t \ll T$, when the walker is mostly located inside $\mathcal{D}_t$ and performs a simple symmetric nearest random walk. f) Survival probability has a function of time for different values of $r$. For a sufficiently large $|\beta|$, the survival probability decays faster than exponentially (we observe a decay consistent with an exponential form), so that $\theta = \infty$.

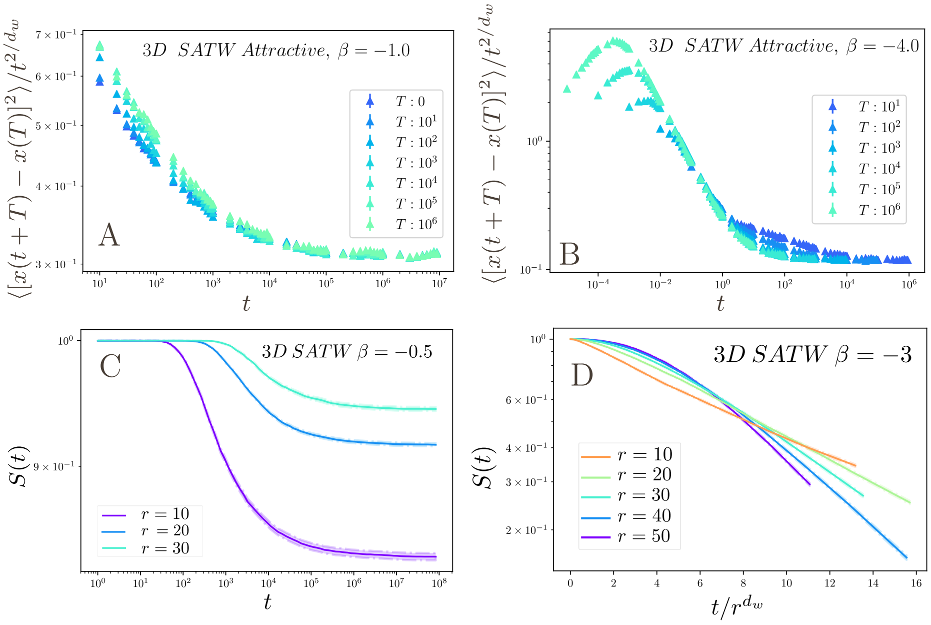


Supplementary Figure 14. First passage properties and aging of the SATW in dimension 3. a) b) Aging of the increments for the SATW (normalized by the expected diffusive scaling at long times). Each curve corresponds to a fixed value of $T$. The increments are stationary at long times at low $|\beta|$ for the diffusive regime (a) contrary to the subdiffusive case (b) where a diffusive regime is recovered for $t \ll T$ . c) d) Survival probability has a function of time (log scale) for different values of $r$. In the diffusive case c), the survival probability tends to a non-zero constant which depends on $r$ : exploration is thus non compact. Conversely for a sufficiently large $|\beta|$, in the subdiffusive case d), the process performs a compact exploration with an exponential-like decay of the survival probability.
